# Non-selective response inhibition in equiprobable Go/NoGo task: Bayesian analysis of fMRI data

**DOI:** 10.1101/823625

**Authors:** Ruslan Masharipov, Alexander Korotkov, Svyatoslav Medvedev, Maxim Kireev

**Author notes:** Corresponding author: Maxim Kireev; Address: Academika Pavlova Street 9, St. Petersburg, 197376, Russia.

## Abstract

Response inhibition is typically considered a brain mechanism selectively triggered by particular “inhibitory” stimuli or events. Based on recent research, an alternative non-selective mechanism was proposed by several authors. Presumably, the inhibitory brain activity may be triggered not only by the presentation of “inhibitory” stimuli but also by any imperative stimuli, including Go stimuli, when the context is uncertain. Earlier support for this notion was mainly based on the absence of a significant difference between neural activity evoked by equiprobable Go and NoGo stimuli. Equiprobable Go/NoGo design with a simple response time task limits potential confounds between response inhibition and accompanying cognitive processes while not preventing prepotent automaticity. However, previous neuroimaging studies utilized classical null hypothesis significance testing, making it impossible to accept the null hypothesis. Therefore, the current research aimed to provide evidence for practical equivalence of neuronal activity in Go and NoGo trials using Bayesian analysis of functional magnetic resonance imaging (fMRI) data. Thirty-four healthy participants performed a cued Go/NoGo task with an equiprobable presentation of Go and NoGo stimuli. To independently localize brain areas associated with response inhibition in similar experimental conditions, we performed a meta-analysis of fMRI studies using equal probability Go/NoGo tasks. As a result, we observed overlap between response inhibition areas and areas demonstrating the practical equivalence of neuronal activity located in the right dorsolateral prefrontal cortex, parietal cortex, premotor cortex, and left inferior frontal gyrus. Thus, obtained results favour the existence of non-selective response inhibition, which can act in settings of contextual uncertainty induced by the equal probability of Go and NoGo stimuli.

**Highlights:**

- Non-selective response inhibition was assessed by equiprobable Go/NoGo task
- Bayesian analysis of fMRI data was combined with a meta-analysis of fMRI studies
- Several nodes of response inhibition system were equally involved in Go and NoGo trials
- Evidence for non-selective response inhibition in uncertain context was found

## 1. Introduction

Response inhibition is the ability to suppress inappropriate, automatic, reflexive, or habitual prepotent responses to produce a controlled goal-directed response (Friedman et al., 2008, Isoda & Hikosaka, 2011). It is generally accepted in the literature that response inhibition works in close relation to other processes associated with cognitive control, such as working memory, voluntary attention, conflict monitoring, and action selection (Buchsbaum et al., 2005, Nee, Wager & Jonides, 2007, Simmonds, Pekar & Mostofsky, 2008, Criaud & Boulinguez, 2013, Cieslik et al., 2015). Moreover, response inhibition is thought to represent a multifaceted phenomenon rather than a unitary brain mechanism. A distinction is made between action cancellation and action restraint (Swick, Ashley & Turken, 2011, Cieslik et al., 2015, Zhang, Geng & Lee, 2017) as well as between reactive and proactive response inhibition mechanisms (Jaffard et al., 2007, 2008, Aron, 2011, Braver, 2012, Erika-Florence, Leech & Hampshire, 2013).

According to the conventional view, the response inhibition process is selectively triggered by “inhibitory” stimuli that result in increased neuronal activity in brain structures responsible for inhibitory control (Verbruggen & Logan, 2008, Verbruggen et al., 2014, Logan et al., 2014, Criaud et al., 2017). However, in several cases, the concept of selective response inhibition fails to explain observed behavioural and neurophysiological phenomena. Manipulating the probability of the occurrence of “inhibitory” stimuli and the subjects’ awareness of the probability of the appearance may slow down the motor response (Chevrier, Noseworthy & Schachar, 2007, Jaffard et al., 2007, Boulinguez et al., 2008, Vink et al., 2014, Vink et al., 2015, Dunovan et al., 2015, Meffert et al., 2016, Hsieh, Wu & Tang, 2016). Moreover, when necessary to rapidly suppress a specific action, the action is inhibited along with all other potential actions. That is, such inhibition can affect the entire motor system (Coxon, Stinear & Byblow, 2007, Aron & Verbruggen, 2008, Badry et al., 2009, Duque & Ivry, 2009, Duque et al., 2010, Majid et al., 2012, MacDonald et al., 2017). In attempting to explain the effects mentioned above, several authors proposed the concept of non-selective (“global”) response inhibition (Frank, 2006, Aron, 2011, Albares et al., 2014, Criaud et al., 2017). It is thought that those non-selective mechanisms serve to prevent inappropriate or premature responses at the expense of the speed of execution of correct actions. First, response inhibition mechanisms may non-selectively inhibit all potential responses to further selectively execute an appropriate response (non-selective inhibition of multiple concurrent motor responses). Second, inhibition may be triggered not only by the presentation of “inhibitory” stimuli but also by the occurrence of any imperative stimuli instructing on the necessity to suppress or execute a prepared action (non-selectivity of inhibitory stimulus perception). In the present work, we consider the latter mechanism.

A tentative neurophysiological model of the non-selective or “global” response inhibition involved in the resolution of interference between several competing response options was proposed by Frank (2006) and included the cortico-subthalamic “hyper-direct” pathway (Nambu et al., 1996, Haynes & Haber, 2013) which is capable of rapidly and non-selectively suppress all potential response options. As the research area developed, it was hypothesized that the model might be applied not only to tasks with multiple concurrent response options but also to simpler tasks where the subject has to choose between executing and refraining from an action (Albares et al., 2014, Criaud et al., 2017). The authors used a cued equiprobable Go/NoGo task, wherein a preparatory cue stimulus indicated the probability of a NoGo stimulus occurrence. A simple equiprobable Go/NoGo was chosen instead of a complex Go/NoGo task to limit confounds between response inhibition and accompanying cognitive processes (Criaud et. al., 2013). Complex Go/NoGo tasks usually utilize the low probability of NoGo stimulus, difficulties in identifying NoGo signals, high attentional or working memory loads. Although one of the possible ways to build up a prepotent response tendency is biasing Go/NoGo probabilities in favour of Go stimuli, it is not necessary when the design involves a simple speeded reaction time task with a single response and reduces the complexity of identification of Go and NoGo signals to one bit of information (Criaud et al., 2017). In these conditions, a stimulus that does not require a response elicits subthreshold automatic motor activations that do not become overt because they are counteracted by fast, automatic response inhibition (Boulinguez et al., 2008, 2009; Sumner & Husain, 2008; McBride et al., 2012; Albares et al., 2015).

Within the framework of the selective inhibition model, the inhibition process would only be triggered by the identification of the NoGo stimulus. According to the hypothesis of non-selective inhibition, when the context is uncertain (equal probability of NoGo and Go stimuli), the need for response inhibition arises for both NoGo and Go trials. Experimental assessment of the hypotheses revealed no statistically significant differences between NoGo and Go trials both in the amplitude of early components of event-related potentials (ERP) (Albares et al., 2014) and in the level of neuronal activity measured by functional magnetic resonance imaging (fMRI) (Criaud et al., 2017). At the same time, a significant difference in early ERP amplitudes (peaked at 170 ms) was found in uncertain equiprobable NoGo and Go trials compared to certain Go-control trials, where no inhibition is required (Albares et al., 2014). The authors considered this fact as evidence favouring the presence of a “non-selective” inhibitory mechanism that is not specific to the processing of NoGo signals. The analysis of early ERP components suggests that non-selective inhibition may blindly suppress any automatic response when the context is uncertain, acting as a gating mechanism controlling the initiation of a prepared, prepotent response.

However, a critical limitation of the abovementioned studies was that the authors could not accept the null hypothesis, so they did not provide direct proof of the practical equivalence of neuronal activity level between Go and NoGo trials in the inhibition-related brain structures. Indeed, within the framework of classical (frequentist) null hypothesis significance testing (NHST), we cannot accept the null hypothesis based on the absence of a significant difference. We can only reject it. Thus, the question of experimental support for non-selective response inhibition remains unanswered, and answering this question requires overcoming the methodological limitation of NHST, which is possible using Bayesian statistics (Kruschke & Liddell, 2017).

Therefore, the present study aimed to verify the non-selective response inhibition hypothesis by using the fMRI data from an equiprobable Go/NoGo task. Based on the results of previous studies, it may be suggested that if the hypothesis on the non-selectivity of inhibition is correct, then the brain structures responsible for response inhibition will demonstrate practically equivalent levels of neuronal activity in equiprobable Go and NoGo trials. Bayesian parameter inference was applied to assess this prediction. Instead, if the hypothesis on the selectivity of response inhibition is correct, then activation of the inhibition-related brain structures will be observed in the NoGo conditions compared with Go conditions.

## 2. Material and methods

### 2.1. A meta-analysis of fMRI studies using equal probability Go/NoGo tasks

Given that a practically equivalent level of neuronal activity may be identified not only for response inhibition-related structures but, for example, those related to sensory processing of visual stimuli, working memory, and attention, we conducted a meta-analysis of fMRI studies to independently localize brain structures associated with response inhibition. Studies using equal probability Go/NoGo tasks were selected for the meta-analysis. We searched for studies that, similar to the investigation by Criaud et al. (2017), compared the neuronal activity in the condition of equiprobable Go and NoGo stimuli presentation with the control Go condition, wherein the subject did not need to inhibit the prepared action.

We searched for these studies in the PubMed database in the period of 01/01/2000 to 15/07/2019 using the following keywords: “((fmri) OR (functional magnetic resonance)) AND ((nogo) OR (no-go)).” Four additional studies were identified through manual searches. As a result, 726 papers were identified. At the first stage of selection, we excluded reviews, meta-analyses, and papers repeatedly reporting the results of fMRI studies (see the flow chart of analysis at Fig. 1).

**Figure 1.**
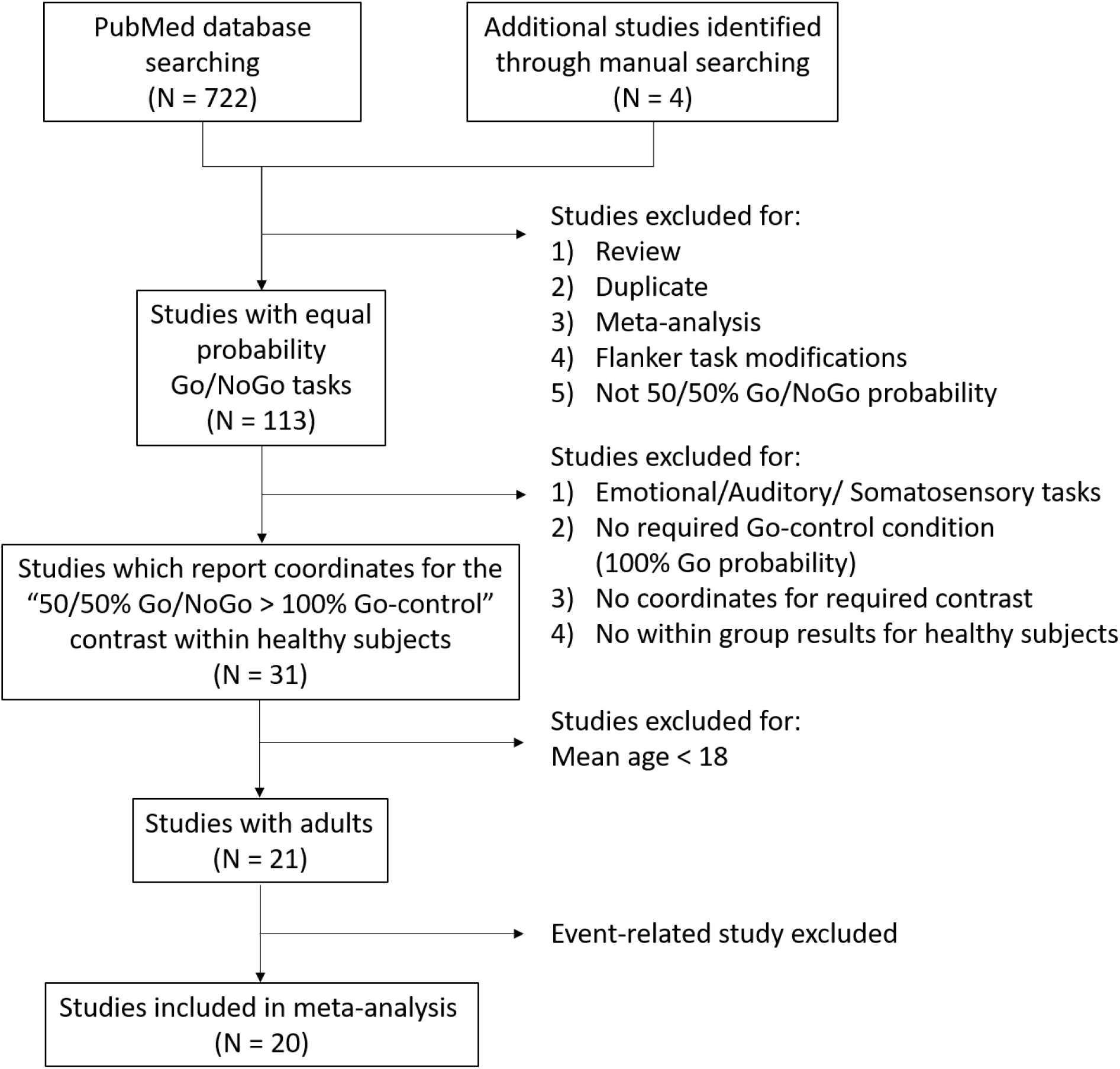
Flow chart of study selection in the meta-analysis.

Studies dealing with Flanker task modifications of Go/NoGo task or based on unequal probability of appearance of Go and NoGo trials were also excluded from the analysis. As a result, 593 papers were excluded. Task designs of the remaining 113 papers provided for equal probability Go and NoGo stimuli presentation. At the next step, 82 of the papers were excluded based on the following criteria: auditory and sensorimotor Go/NoGo tasks were used; only emotion-laden task conditions were used (emotionally neutral conditions were either absent or not considered separately); required Go-control condition (100% probability of the Go stimulus presentation) was not used; the coordinates for the contrast of interest “50/50% Go/NoGo > 100% Go-control” within a group of healthy subjects were not reported. In 11 out of the 31 remaining studies, healthy volunteer subjects under the age of 18 (children and adolescents) were studied. These papers were also excluded from analysis because this study focused on the brain activity of healthy, adult subjects. In all remaining studies except one, only block designs were used. To make our sample more homogenous, we excluded the only eligible study with an event-related design (Criaud et al., 2017). The final meta-analysis included 20 studies (452 healthy subjects, mean age of 29 years) with a total of 210 foci (for more details, see supplementary materials, Table S1).

Coordinate-based meta-analysis was performed using the random-effects activation likelihood estimation (ALE) algorithm (Eickhoff et al., 2009, 2012; Turkeltaub et al., 2002, 2012) implemented in GingerALE 3.0.2 software (http://brainmap.org/ale). The ALE algorithm assesses the spatial convergence between neuroimaging studies by modelling spatial uncertainty of activation foci using three-dimensional Gaussian probability functions with the full-width at half-maximum (FWHM) inversely related to the square root of the sample size of the original study (Eickhoff et al., 2009). An ALE map is obtained by computing the union of activation probabilities across studies for each voxel and tested against a null-distribution of random spatial convergence between studies (Eickhoff et al., 2012). Statistical significance was determined using a cluster-level threshold of 0.05 corrected for family-wise error (FWE) with an uncorrected cluster-forming threshold of 0.001 (5000 thresholding permutations), which was observed to provide the best compromise between sensitivity and specificity for ALE analysis (Eickhoff et al., 2016). All the coordinates were converted into Montreal Neurological Institute (MNI) space using the Lancaster transform (Lancaster et al., 2007). An additional FWHM of 4 mm was used so that the median FWHM value of Gaussian functions was at 13.5 mm to match the estimated spatial smoothness of the preprocessed fMRI data to be analysed for conjunction in the current study. The number of studies included in the current meta-analysis meets the minimum recommended number of 17–20 studies for the ALE algorithm (Eickhoff et al., 2016).

### 2.2. Subjects

The recruitment of subjects for the present fMRI study was carried out in two stages. At the first stage, the sample size was 20 subjects (16 women, aged (mean ± SD) 23.9 ± 4.6). During the review process, the concern regarding the potential impact of relatively small sample size on the observed null-effects was raised. As a result, a retrospective power analysis was carried out which suggested increasing the sample size up to 34 subject (see section 2.3). Therefore, the final sample consisted of 34 healthy, right-handed volunteer subjects (24 women, aged 25.9 ± 5.2). An Oldfield test was used to determine the dominant arm (Oldfield, 1971). The subject volunteers signed written informed consent to participate in the study and were paid for their participation. All the procedures were performed in accordance with the Declaration of Helsinki and were approved by the Ethics Committee of the N.P. Bechtereva Institute of the Human brain of the Russian Academy of Sciences.

### 2.3. Power analysis

Within the classical NHST framework, the question may arise as to whether the sample size was sufficient to detect selective inhibition when a non-significant result was obtained. To determine a sufficient sample size, a prospective power analysis is universally accepted while the informativeness of the retrospective power is debatable (Thomas, 1997). The “observed power” based on the obtained data (obtained effect size and variance) is clearly uninformative, as it is directly related to the p-value (Hoenig & Heisey, 2001). The other two types of retrospective power analysis are based on pre-specified parameters (predicted effect size *or* variance) or independent datasets (predicted effect size *and* variance) (Nakagawa & Foster, 2004). The latter one breaks the circularity of the “observed power” and sometimes can potentially be useful. (Thomas, 1997; O’Keefe, 2007). To perform power analysis, we used the Consortium for Neuropsychiatric Phenomic LA5c dataset (Poldrack et al., 2016, Gorgolewski et al., 2017) as it contains fMRI data from a relatively large cohort (N = 115) of healthy subjects performing a stop-signal task (see supplementary materials for details). This classical inhibitory paradigm models a situation where the inhibitory brain activity is selectively triggered by infrequent inhibitory stimuli (stop signal) but not by Go stimuli (Criaud et al., 2017). The mean effect sizes for the selective inhibition contrast (“Correct-Stop > Go” contrast, see Fig. S1) were estimated within the brain regions revealed by the current meta-analysis. They ranged from 0.57 to 1.12 Cohen’s d. Power analysis was performed using GPower 3.1.9.7 (Faul et al., 2007). It indicated that the sample size of 34 subjects would be sufficient to detect the selective inhibition effect (d = 0.57, two-tailed one-sample test, alpha = 0.05) with the power of 0.9 (see Fig. S2). The power of 0.8 or 0.9 is commonly used for sample size calculations (Dell, Holleran & Ramakrishnan, 2002).

However, it is worth noting that obtaining non-significant results with high retrospective power (based on an independent dataset) or high prospective power would not provide direct evidence for the null hypothesis (Denies and Mclatchie, 2018). To provide evidence for the null hypothesis Bayesian inference or frequentist equivalence testing should be used (Kruschke & Liddel, 2015, Lakens, 2017, Kruschke, 2018). Accordingly, in the current study, all conclusions were made only on the basis of the Bayesian analysis and its overlap with the meta-analysis. Results for the classical NHST was added only to illustrate how the more familiar frequentist inference relates to Bayesian inference.

### 2.4. Experimental task and study procedure

We used a paired stimulus modification of the Go/NoGo task (see Fig. 2) (Kropotov, 2016). This task was originally developed to dissociate cognitive processes, such as response inhibition, conflict monitoring, sensory mismatch and category discrimination (Kropotov et al., 2011, 2015). Given that imperative Go and NoGo stimuli were presented in an equiprobable manner this modification of the Go/NoGo task was used for the purposes of the current study. Each trial consisted of the two consequently presented stimuli. The first preparatory cue stimulus warned the subjects on the presentation of the second, imperative stimulus, or indicated no need for any response to the second stimulus. The study included two variants of the experiment’s instructions. According to the first instruction (see Fig. 2A), the subject should press the response button as soon as possible upon presentation of the pair of images “animal-animal” (“A-A Go” trials) and refrain from acting upon presentation of the pair “animal-plant” (“A-P NoGo” trials). According to the second experiment’s instruction (see Fig. 2B), the subject acts after the presentation of the pair “animal-plant” (“A-P Go” trials) and suppresses an action upon presentation of the pair “animal-animal” (“A-A NoGo” trials). The subjects were familiarized with the task just before scanning to ensure they understood the instructions. Additionally, before each fMRI session, the subjects were reminded of the need to react to Go stimuli as quickly as possible while refraining from reacting to NoGo stimuli.

**Figure 2.**
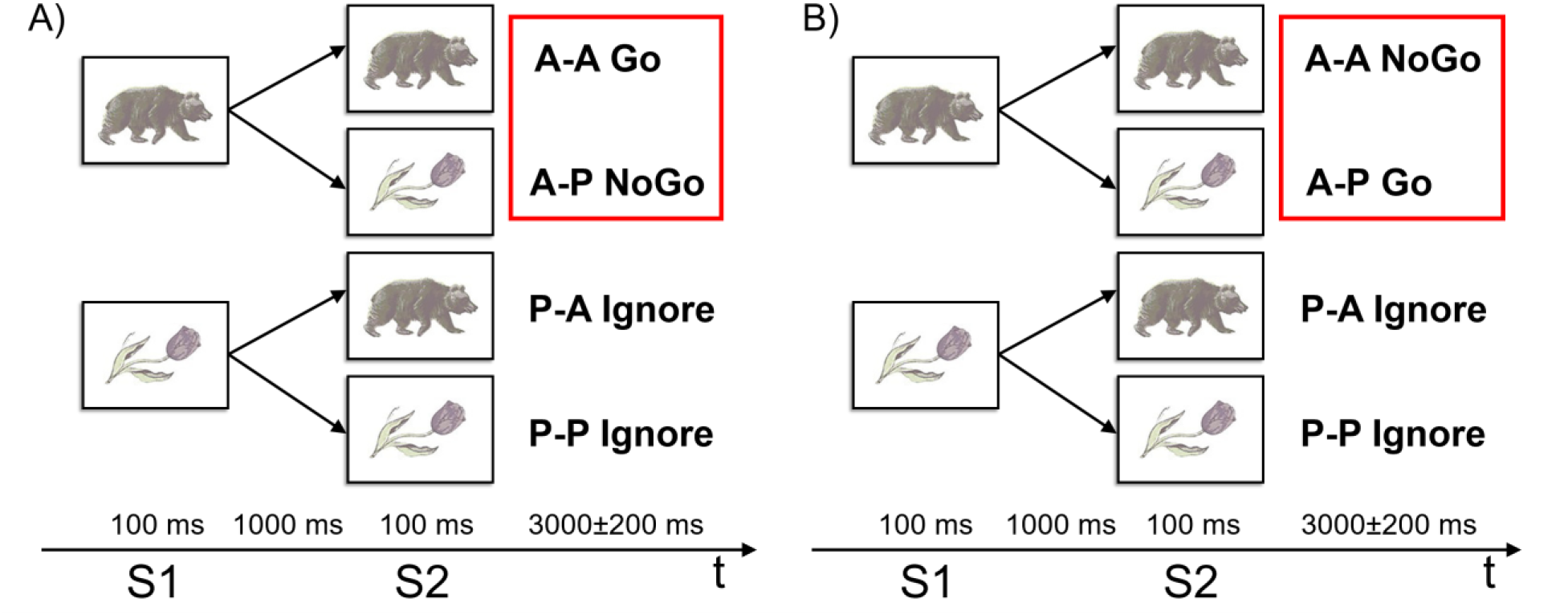
Experimental design of the Go/NoGo task. S1 – first stimulus (preparatory cue). S2 – second stimulus (imperative stimulus). A – images of animals, P – images of plants. A) First experiment: “A-A Go”, “A-P NoGo”. B) Second experiment: “A-P Go”, “A-A NoGo”. Red boxes highlight trials that were compared to test the hypothesis of selective and non-selective response inhibition.

In both experiments, if the first presented stimulus was an image of a “plant,” subjects do not need to perform any actions in response to the presentation of any second stimuli of a trial. In such a trial, the subject should ignore the second stimulus and wait for the next pair of stimuli (“P-A Ignore” and “P-P Ignore” trials). Accordingly, it was assumed that there is no inhibition of prepared action in “Ignore” conditions. The “Ignore” trials were included in the adopted Go/NoGo task to evaluate preparatory brain activity, such as preparing to receive a relevant stimulus (attentional set) and preparing to make a movement (motor set), by comparing them to Go and NoGo trials. However, the issue related to preparatory processes is beyond the scope of the present study. Fifty pairs of each type of stimuli were randomly presented in each experiment. The order of following the instructions was counterbalanced among the subjects. Two variants of the present Go/NoGo task made it possible to control the differences in the load on working memory between A-P/A-A - Go/NoGo stimuli (Kropotov & Ponomarev, 2015).

The fMRI data obtained using this experimental task allowed us to test the hypothesis on the *non-selectivity* of response inhibition in the current study, given that a similar task design was previously used for the same goal by Criaud et al. (2017). The imperative Go and NoGo stimuli were presented after the preparatory stimulus with equal probability, as reported in event-related studies by (Albares et al., 2014, Di Russo et al., 2016, Kropotov, 2016, Criaud et al., 2017). To build up prepotent tendency to react, subjects were instructed to press the button by the right thumb as quickly as possible. This is a case of a simple speeded reaction time task with a single response and only one bit of information. It is known that these conditions elicit subthreshold automatic motor activations (Criaud et al., 2017). According to G.A. Miller: “One bit of information is the amount of information that we need to make a decision between two equally likely alternatives” (Miller, 1956). Here it refers to discrimination between “animal” and “plant” stimuli.

Such an equally probable presentation of imperative stimuli provides several advantages. First, it minimizes the difference in cognitive load between Go and NoGo conditions arising from task complexity (Criaud & Boulinguez, 2013, Criaud et al., 2017). Second, such a design enables the exclusion of effects associated with a low frequency of the NoGo stimuli presentation (“oddball” effects) confounding the effect of inhibition (Di Russo et al., 2016). Third, it creates maximum uncertainty regarding the probability of the presentation of an imperative stimulus and thus minimizes the conflict between two response types, which makes it possible to distinguish between error monitoring and conflict resolution processes on the one hand and response inhibition processes on the other (Lavric, Pizzagalli & Forstmeier, 2004).

In total, 100 NoGo trials, 100 Go trials, 100 P-A Ignore trials, and 100 P-P Ignore trials were presented over two experimental sessions. In the absence of stimulation, a fixation cross was displayed in the centre of the screen. The stimuli were presented for 100 ms, and the interstimulus interval was 1000 ms. The intertrial interval jittered from 2800 to 3200 ms with an increment step of 100 ms. Additionally, to improve design efficiency, 100 zero-events (fixation crosses) were randomly inserted between the stimuli pairs (trials), and their duration jittered from 3000 to 5000 ms with an increment size of 500 ms. The action to be performed consisted of pressing a button with the right thumb. The duration of one task session was 17.5 minutes. Before starting the fMRI study, the subjects performed a training task. The Invivo’s Eloquence fMRI System (Invivo, Orlando, FL, USA) was used to deliver the stimuli, synchronize with fMRI acquisition, and record reaction times of subjects bottom pressing. The task presentation sequence and all temporal parameters of the stimuli presentation were programmed using the E-prime 2.0 software package (Psychology Software Tools Inc., Pittsburgh, PA, USA).

### 2.5. Image acquisition

A Philips Achieva 3.0 Tesla scanner (Philips Medical Systems, Best, Netherlands) was used for the study. The structural T1-images were registered with the following parameters: field of view (FOV) – 240×240 mm, repetition time (TR) – 25 ms, echo time (TE) – 2.2 ms, 130 axial slices with a thickness of 1 mm and pixel size of 1×1 mm, flip angle – 30°. For the registration of T2*-images, a single-pulse echo planar imaging (EPI) sequence was used. The period of data registration from 31 axial slices was 2 seconds (TR = 2 s, TE = 35 ms). The following parameters were employed: FOV – 200×186 mm, flip angle – 90°, voxel size – 3×3×3 mm. Two dummy scans were performed prior to each session. To minimize head motions, we used an MR-compatible soft cervical collar.

### 2.6. Preprocessing of fMRI-images

Image preprocessing included the following: realignment to the first image of the session, slice time correction, co-registration, segmentation, normalization to an MNI template, and spatial smoothing (8 mm FWHM). Preprocessing and statistical analyses of the images were performed using an SPM12 (Statistical parametric mapping) software package (http://www.fil.ion.ucl.ac.uk/spm).

### 2.7. Statistical analysis of fMRI data

The first level of analysis was conducted using frequentist parameter estimation. The second level of analysis was conducted using both the frequentist and Bayesian parameter estimation (Friston et al., 2002a, Friston & Penny, 2003, Neumann & Lohmann, 2003). Onset times of second stimuli presentation (separately for “A-A Go”, “A-P NoGo”, “P-A Ignore Exp1”, “P-P Ignore Exp1”, and also “A-P Go”, “A-A NoGo”, “P-A Ignore Exp2”, “P-P Ignore Exp2”), erroneous button pressing, and missing the responding in Go trials were used to create regressors of the general linear model (GLM) for each subject. Events were impulse responses with a duration of zero convolved with the canonical haemodynamic response function (HRF). Low-frequency drift was removed by temporal high pass filtering with a cut-off frequency of 1/128 Hz. Six head motion parameters were included in the GLM as nuisance regressors to account for the movement artefacts (Johnstone et al., 2006). Beta-coefficients reflecting an increase in the blood oxygenation level-dependent (BOLD) signal in the experimental condition relative to the implicit baseline were scaled to percent signal change (PSC) following the procedure recommended in (Poldrack, Mumford, Nichols, 2011, p.186). To that end, beta-coefficients for each condition were divided by the mean value of beta-coefficients for the constant term and multiplied by 100 and a scaling factor (SF) that is needed so that the peak of an isolated BOLD response is equal to one (SF = 0.21). Two linear contrasts of scaled beta-coefficients were calculated: (1) 0.5 × [“A-P NoGo” + “A-A NoGo”] – 0.5 × [“A-P Go” + “A-A Go”] (*the “NoGo vs. Go” comparison*); (2) 0.25 × [“A-P NoGo” + “A-A NoGo” + “A-P Go” + “A-A Go”] – 0.25 × [“P-A Ignore Exp1” + “P-P Ignore Exp1” + “P-A Ignore Exp2” + “P-P Ignore Exp2”] (*the “Go + NoGo vs. Ignore” comparison*). The sum of positive contrast weights was equal to one. The contrasts from 34 subjects were used as variables to verify the hypotheses on selective and non-selective response inhibition at the second level of analysis. Only grey-matter voxels were included in the second-level analysis. To that end, a mask was created based on the segmentation of each subject’s structural T1-images.

### 2.8. Verification of the hypotheses on selective and non-selective response inhibition

As in previous studies employing equiprobable Go/NoGo task with a single prepotent motor response, in the current study, non-selectivity refers to the perceptual decision mechanisms involved in the detection, discrimination, or identification of sensory stimuli (Albares et al., 2014; Criaud et al., 2017). In particular, we did not consider non-selectivity related to the decision mechanisms that involve the selection between multiple alternative responses, which confound response inhibition processes in choice reaction time tasks (Criaud et al., 2013, 2017). The former type of non-selectivity implies that any imperative stimuli (both Go and NoGo stimuli) would trigger response inhibition when the context is uncertain because of equal probability of Go and NoGo stimuli. The later type of non-selectivity, usually called “global” inhibition, implies that inhibition affects all alternative responses included the selected response (Aron & Verbruggen, 2008).

We tested the hypotheses on selective and non-selective response inhibition using the “NoGo vs. Go” comparison. In the case of the *selectivity* of response inhibition, we expect to find a selective increase in the neuronal activity in response to the presentation of NoGo stimuli compared to Go stimuli (“NoGo > Go”). In the case of the *non-selectivity* of response inhibition, we expect to find a practically equivalent increase in the neuronal activity in response to the presentation of both NoGo and Go stimuli (“NoGo = Go”) in the brain areas related to the response inhibition, which were independently localized by the current meta-analysis (“50/50% Go/NoGo blocks > 100% Go-control blocks”). Additionally, we used the “Go + NoGo vs. Ignore” comparison to distinguish between the brain areas that are simply not activated in current task settings (“Go + NoGo = Ignore”) from the brain areas activated in equiprobable Go and NoGo trials compared to Ignore trials, where no inhibition is required (“Go + NoGo > Ignore”). Thus, in order to identify the non-selective response inhibition in the settings of contextual uncertainty induced by the equal probability of Go and NoGo stimuli, it was necessary to show three-way overlap between (1) inhibitory-related brain areas according to the meta-analysis (“50/50% Go/NoGo blocks > 100% Go-control blocks”), (2) brain areas with practically equivalent neuronal activity in Go and NoGo trials (“NoGo = Go”) and (3) brain areas activated in equiprobable Go and NoGo trials (“Go + NoGo > Ignore”). This conjunction analysis was performed by binarization and multiplication of thresholded images (Nichols et al., 2005).

To reveal brain areas with practically equivalent neuronal activity in Go and NoGo trials (“NoGo = Go”), one has to provide evidence for the null hypothesis. The classical NHST approach estimates the probability (p-value) of obtaining actual data’s best-fitting parameter value, *β*, or something more extreme under the null hypothesis that the experimental effect is *θ = cβ = 0*. One can never accept the null hypothesis using NHST because the p-value does not represent the probability of the null hypothesis, and the probability an effect equals exactly zero is itself zero (Friston et al., 2002a). A non-significant result can be obtained in two cases (Dienes, 2014): (1) there is no effect, and our data are against the alternative hypothesis; or (2) our data are insufficient to distinguish alternative hypothesis from the null hypothesis, and we cannot confidently make any inference (low statistical power). When the obtained result is not significant, it is recommended to use frequentist equivalence testing (Lakens, 2017) or Bayesian inference (Penny & Ridgway, 2013; Dienes, 2014; Kruschke & Liddell, 2017, Kruschke, 2018) to provide evidence for the null hypothesis. In the present study, the Bayesian parameter inference was utilized (Friston & Penny, 2003; Penny & Ridgway, 2013; Kruschke, 2018).

The Bayesian approach estimates the posterior probability (*PP*) distribution of the effect *θ*, given the data using the likelihood and the prior knowledge. For the second-level Bayesian analysis, SPM12 implements the hierarchical parametric empirical Bayes approach with the global shrinkage prior (Friston & Penny, 2003). It represents a prior belief that, on average, in the whole brain, there is no global experimental effect. If the posterior probability of the effect exceeding the effect size threshold, *γ*, is greater than the predefined probability threshold, *α* = 95%, then the hypothesis on the presence of “NoGo > Go” effect will be accepted (see Fig. S3A in the supplementary materials):

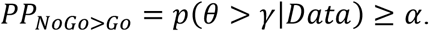

If the effect value falls within the interval [−*γ*; *γ*] with a probability of *α* = 0.95, then the hypothesis of the null “NoGo = Go” effect will be accepted, supporting the practical equivalence (Kruschke, 2018) of the BOLD signal between compared conditions (see Fig. S3B):

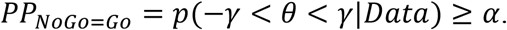

The interval [−*γ*; *γ*] can be thought of as the neuronal “background noise level” (Eickhoff et al., 2008) or as a region of practical equivalence (ROPE) that expresses which effect size values (PSC) are equivalent to the null value for current practical purposes (Kruschke & Liddell, 2017). The hypothesis on the presence of the “Go > NoGo” effect will be accepted if (see Fig. S3C):

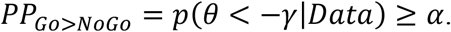

If none of the above criteria are satisfied, the data in particular voxel are insufficient to distinguish the null hypothesis from the alternative hypothesis (“low-confidence” voxels, see Fig. S3D) (Magerkurth et al., 2015). For such voxels, it is impossible to make any inference confidently, so we need to increase the sample size or scanning time. In the current study, our conclusions were based only on voxels in which the effect size values (PSC) fell within or outside the interval [−*γ*; *γ*] with the *PP* > 0.95. Thus, for these voxels, the obtained data are sufficient to make an inference confidently. This decision rule is also known as the “ROPE-only” decision rule (Kruschke, 2018, see Fig. S3). It was applied for the “NoGo vs. Go” and “Go + NoGo vs. Ignore” comparisons. Bayesian parameter inference was performed using in-house scripts based on SPM12 (https://github.com/Masharipov/Non-selective-inhibition). Bayesian parameter inference based on the “ROPE-only” decision rule with a zero effect size threshold corresponds to the false discovery rate correction for multiple comparisons within the NHST framework (Genovese, Lazar & Nichols, 2002, Friston & Penny, 2003). The group-level effect size threshold *γ* was set at one standard deviation of the prior variance of the contrast (prior SD_*θ*_), which is the default in SPM12 (Friston & Penny, 2003; Eickhoff et al., 2008). For visualization purposes, posterior probabilities were converted to the log posterior odds, LogOdds = log(*PP*/(1-*PP*)). LogOdds > 3 correspond to *PP* > 0.95. Anatomical localization of clusters was identified using xjView toolbox (http://www.alivelearn.net/xjview).

### 2.9. Sequential analyses

Since the statistical analysis was carried out twice, for the sample size N = 20 and N = 34, the question of “data peeking” should be addressed. The “data peeking” refers to the practice when a researcher periodically reanalyzes sequentially obtained data and decides to stop data collection as soon as significant results are obtained (Yarkoni & Braver, 2010). The sequential analyses and optional stopping in the classical NHST framework substantially inflate the number of false positives and require special adjustments of p-values along with the definition of the stopping rule based on the power analysis (Lakens, 2014). In contrast, Bayesian inference based on the posterior probabilities or Bayes factors does not depend on stopping intentions and does not necessarily require power analysis (Wagenmakers, 2007; Kruschke and Liddell, 2017). Therefore, the issue of “data peeking” may be mitigated by the usage of Bayesian statistics (Hartshorne and Adena Schachner, 2012).

Within the classical NHST framework, a statistically significant difference may be shown in any voxel with a sufficiently large sample size, even when the effect is trivial or has no practical significance (Friston et al., 2002a). Thus, with an unbounded increase of the data, the number of statistically significant voxels will approach 100% of the total number of voxels (see, for example, Gonzalez-Castillo et al., 2012). At the same time, Bayesian inference allows one to find not only “activated” voxels but also “null effect” voxels with a trivial difference (practical equivalence) and “low confidence” voxels for which obtained data are not sufficient. Increasing sample size using Bayesian parameter inference will lead to the situation when the number of “activated” and “null effect” voxels reaches a plateau. When the plateau was reached, it is reasonable to stop the data collection. To illustrate how this relates to the current study we provided analysis of the dependencies between the sample size and the number of “activated”, “null effect,” and “low confidence” voxels for the “NoGo vs. Go” comparison. To plot these dependencies the analysis was performed for the samples from 6 to 34 subjects with a step of 2 subjects (see the supplementary materials). In total thirty random groups were sampled for each step.

## 3. Results

### 3.1. Meta-analysis

As a result of the meta-analysis of 20 fMRI studies, we identified the brain structures that were characterized by increased neuronal activity demonstrated in the settings of equal probability Go and NoGo stimuli presentation compared to the control Go conditions (“50/50% Go/NoGo blocks > 100% Go-control blocks” contrast): (1) right dorsolateral prefrontal cortex (DLPFC), (2) right inferior parietal lobule (IPL), (3) right temporoparietal junction (TPJ), (4) bilateral inferior frontal gyrus (IFG) and anterior insula (also known as anterior insula/frontal operculum (AIFO)), (5) right premotor cortex (PMC) and frontal eye field (FEF), (6) bilateral anterior cingulate cortex (ACC) and supplementary motor area (SMA), and (7) bilateral thalamus (see Fig. 3, Table 1).

**Table 1.** The result of the meta-analysis of 20 fMRI studies using equal probability Go/NoGo tasks (“50/50% Go/NoGo blocks > 100% Go-control blocks” contrast).

| Nº | Cluster size, mm<sup>3</sup> | Peak MNI-coordinate, mm | Peak ALE-score | Anatomical localization (L – left, R – right hemisphere; BA – Brodmann area) |
| --- | --- | --- | --- | --- |
| 1 | 10160 | 42 34 32 | 0.0114 | R: DLPFC (middle frontal gyrus), BA 8,9 |
| 2 | 8368 | 46 18 -10 | 0.0138 | R: IFG/Anterior insula (AIFO), BA 13, 45, 47 |
| 3 | 4896 | -34 24 0 | 0.0088 | L: IFG/Anterior insula (AIFO), BA 13, 44, 45, 47 |
| 4 | 4176 | 60 -44 26 | 0.0090 | R: TPJ (supramarginal gyrus and superior temporal gyrus), BA 22, 39, 40 |
| 5 | 3408 | 5 18 40 | 0.0103 | R/L: SMA, ACC, BA 6, 8, 24, 32 |
| 6 | 1664 | 12 -8 10 | 0.0089 | R: Thalamus |
| 7 | 1584 | 42 6 40 | 0.0076 | R: PMC, FEF (precentral gyrus, middle frontal gyrus), BA 6, 8, 9 |
| 8 | 1456 | -12 -10 10 | 0.0084 | L: Thalamus |
| 9 | 1424 | 30 -62 38 | 0.0071 | R: IPL (angular gyrus), BA 7, 39 |

Abbreviations: DLPFC – dorsolateral prefrontal cortex, IFG – inferior frontal gyrus, AIFO – anterior insula/frontal operculum, TPJ – temporoparietal junction, SMA – supplementary motor area, ACC – anterior cingulate cortex, PMC – premotor cortex, FEF – frontal eye field, IPL – inferior parietal lobule.

**Figure 3.**
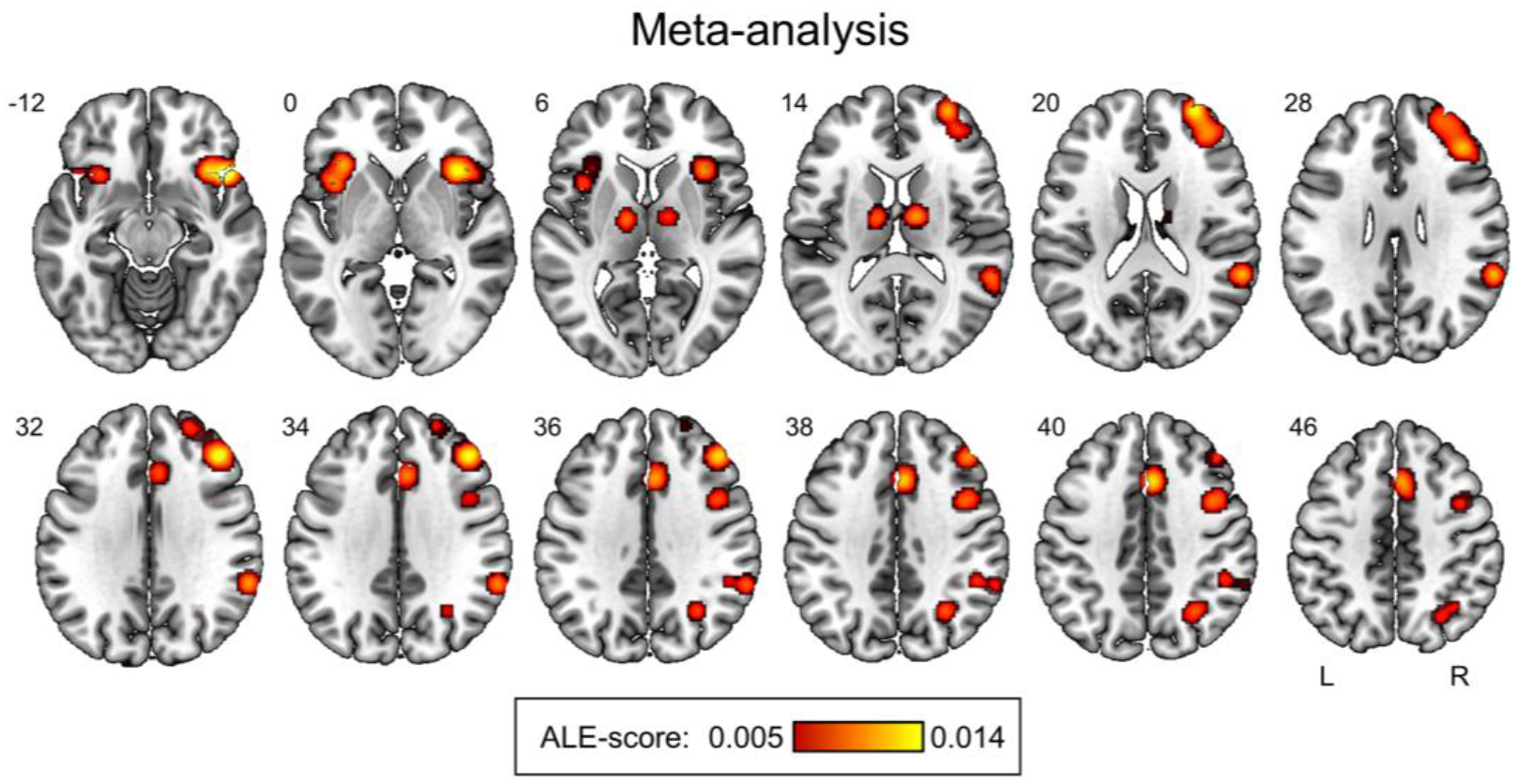
Results of the meta-analysis. The result of the meta-analysis of 20 fMRI studies using equal probability Go/NoGo tasks (“50/50% Go/NoGo blocks > 100% Go-control blocks” contrast).

### 3.2. Behavioural data

In two fMRI sessions, the mean response omission in Go trials was 2.97 ± 3.92%. The mean of false alarms in NoGo trials was 0.35 ± 0.64%. The mean response time (RT) was 384 ± 60 ms, which closely match behavioural data from previous studies with similar equiprobable Go/NoGo design (the mean RT is 380 ms), where subjects were trained to react as fast as possible to create prepotent response tendency (Criaud et al., 2012, 2017; Albares et al., 2014).

### 3.3. fMRI data

Neither classical NHST nor Bayesian inference applied in the present fMRI study revealed a significant increase in neuronal activity in NoGo trials compared to Go trials. Thus, the “NoGo > Go” effect predicted by the hypothesis of selective response inhibition was not observed. The reversed “Go > NoGo” contrast showed the expected motor activations in the pre- and postcentral gyrus, premotor cortex, supplementary motor area, subcortical regions and cerebellum (see Fig. 4 and Tables S2, S3 in the supplementary materials). Both statistical methods revealed similar activation patterns for the “Go > NoGo” effect (Dice coefficient = 0.8), demonstrating the consistency between utilized Bayesian parameter inference with *γ* = 1 prior SD_*θ*_ and LogOdds > 3 (*PP* > 0.95) thresholding approach and NHST with pFWE < 0.05.

**Figure 4.**
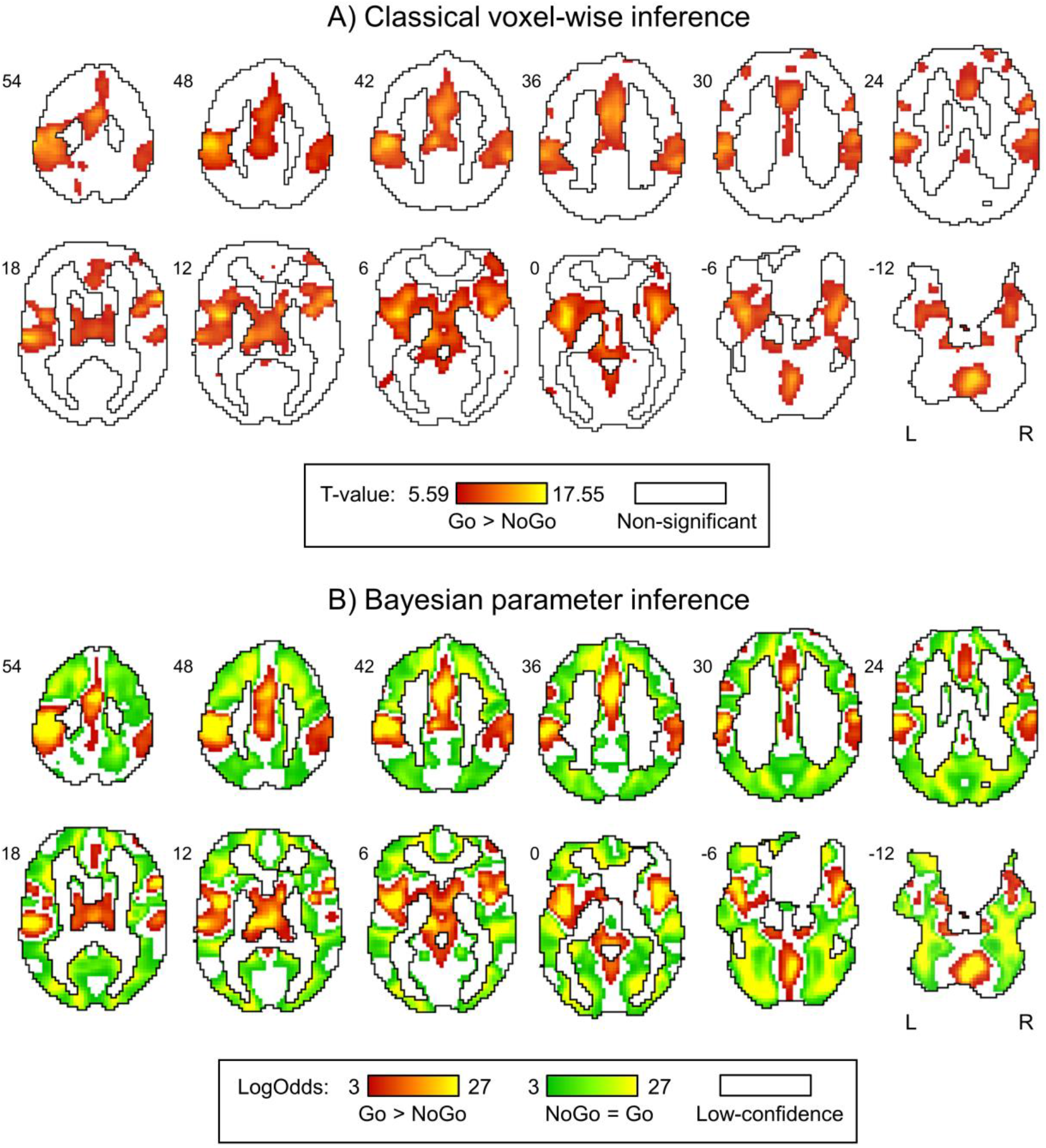
Results of classical voxel-wise and Bayesian parameter inference. A) Classical NHST inference using an FWE-corrected voxel-wise threshold of p < 0.05. B) Bayesian parameter inference using the effect size threshold *γ* = 1 prior SD_*θ*_ = 0.1%, LogOdds > 3 (*PP* > 0.95). The “NoGo > Go” effect was not detected by any of the methods. Red colour depicts the “Go > NoGo” effect. Green colour depicts the “NoGo = Go” effect (practical equivalence of neuronal activity in Go and NoGo trials). White colour depicts A) non-significant voxels and B) “low-confidence” voxels.

At the same time, Bayesian analysis has made it possible to define the brain structures with practically equivalent neuronal activity in Go and NoGo trials. The null “NoGo = Go” effect was revealed for a widely distributed set of regions throughout the whole brain surrounding clusters of activations revealed in the “Go > NoGo” contrast (see Fig. 4A). The “NoGo = Go” regions were separated from activation clusters by regions that consisted of “low-confidence” voxels (Magerkurth et al., 2015). For “low-confidence” voxels, the data obtained are insufficient to accept or reject the null hypothesis. The number of “low-confidence” voxels were decreased with increasing sample size (see Fig. S4 in the supplementary materials). The largest gain in the number of “NoGo = Go” and “Go > NoGo” voxels can be noted from 6 to 20 subjects. After 20 subjects, the dependencies between the sample size and the number of “NoGo = Go” and “Go > NoGo” voxels reached a plateau. This is also confirmed by the similarity between the results of Bayesian parameter inference for the sample size of 20 and 34 subjects (Dice coefficient = 0.88 for the “NoGo = Go” effect, see also Fig. S5 and Table S4 in the supplementary materials).

It should be noted that the “NoGo = Go” area cannot be reasonably split into several parts, such as activation clusters. Therefore, we limited ourselves to specifying the anatomic localization of brain areas where the “NoGo = Go” area overlaps with the clusters revealed by the current meta-analysis and brain areas activated by equiprobable Go and NoGo trials compared to Ignore trials (see results for the “Go + NoGo > Ignore” contrast in the supplementary materials, Fig. S6 and Table S5).

### 3.4. Conjunction analysis

As it can be inferred from the revealed overlap between the results of the meta-analysis and the results of the abovementioned Bayesian analysis, only a few of the brain structures demonstrated (1) increased activity in the “50/50% Go/NoGo blocks > 100% Go-control blocks” meta-analytical comparison, (2) the practical equivalence of the BOLD signal in the “NoGo vs. Go” comparison and (3) activation in Go and NoGo trials compared to Ignore trials. Three-way overlap was observed in (1) right DLPFC, (2) right IPL, (3) right TPJ, (4) left IFG and anterior insula (AIFO), (5) right PMC, and FEF (see Fig. 5, Table 2).

**Table 2.**
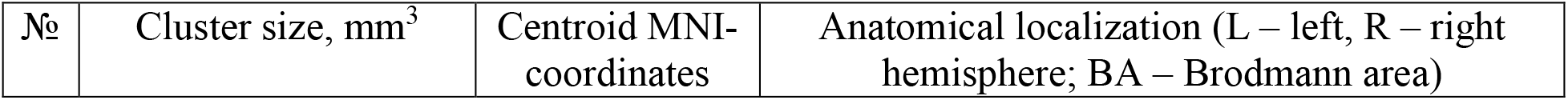
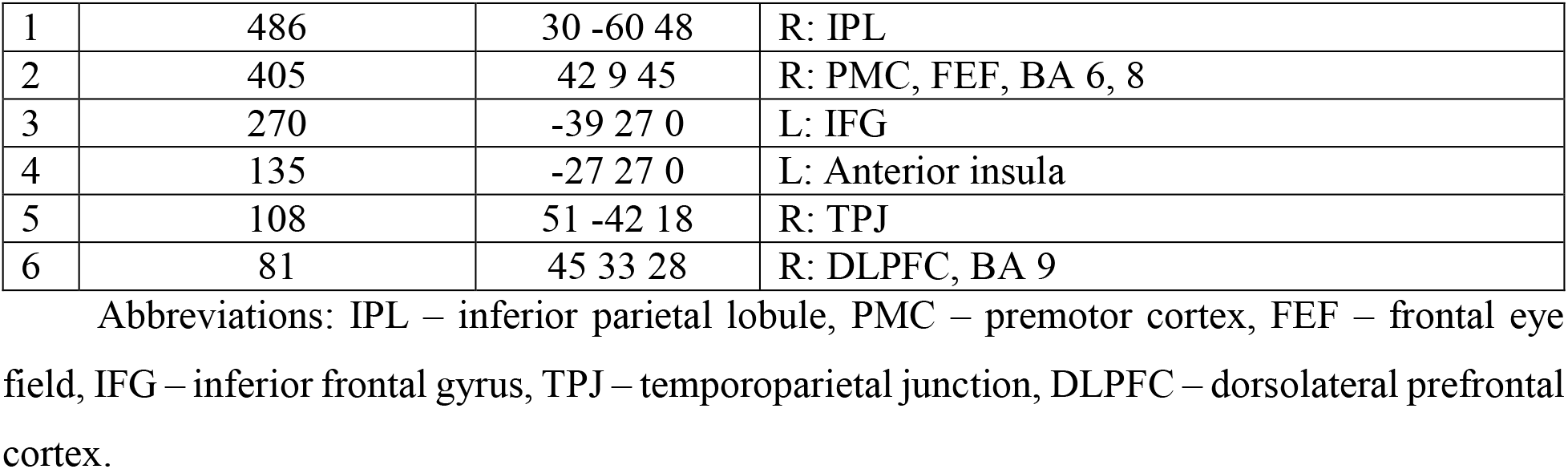
Three-way overlap between the null “NoGo = Go” effect regions revealed by Bayesian analysis of the obtained fMRI data and the results of the current meta-analysis.

**Figure 5.**
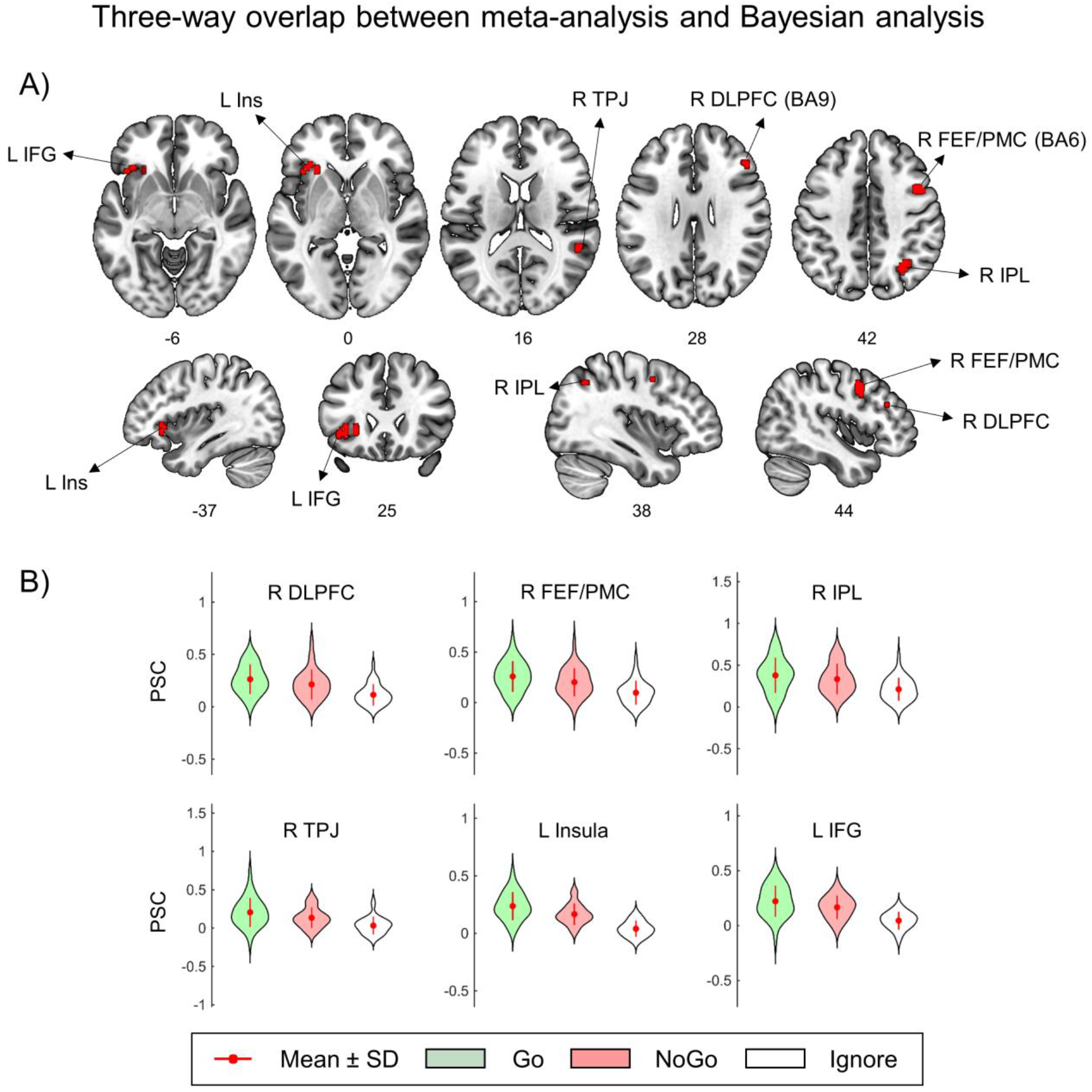
Three-way overlap between meta-analysis and Bayesian analysis. A) Three-way overlap between (1) inhibitory-related brain areas according to the meta-analysis (“50/50% Go/NoGo blocks > 100% Go-control blocks”), (2) brain areas with practically equivalent neuronal activity in Go and NoGo trials (“NoGo = Go”) and (3) brain areas activated in equiprobable Go and NoGo trials (“Go + NoGo > Ignore”). B) Violin plots of the mean PSC in the revealed clusters. Abbreviations: DLPFC – dorsolateral prefrontal cortex, FEF – frontal eye field, PMC – premotor cortex, IPL – inferior parietal lobule, TPJ – temporoparietal junction, Ins – insula, IFG – inferior frontal gyrus, BA – Brodmann area, L – left, R – right.

## 4. Discussion

The results of the present study confirmed the hypothesis of non-selective response inhibition in the contextual uncertainty associated with the equiprobable presentation of Go and NoGo stimuli. Bayesian analysis of the obtained fMRI data provides evidence of the practical equivalence of neuronal activity evoked by Go and NoGo stimuli in several brain structures. According to the results of our meta-analysis, some of these brain structures, including right DLPFC, IPL, and TPJ, left IFG (AIFO), right PMC, and FEF were associated with response inhibition. These structures were repeatedly found activated whenever the condition of equal probability Go and NoGo stimuli presentation was compared with the control Go condition (“50/50% Go/NoGo blocks > 100% Go-control blocks” contrast). In addition, we did not observe the “NoGo > Go” effect in the current study. This finding is consistent with the data of several previous event-related fMRI studies that did not find significant activity increase in NoGo trials compared to Go trials under similar experimental conditions (Lee et al., 2015, Di Russo et al., 2016, Criaud et al., 2017, see also random-effects analysis within visual and audio modalities in Laurens, Kiehl & Liddle, 2004).

However, the question is how precisely observed activity can be associated with response inhibition per se in those brain regions in which the overlap was found between the meta-analysis and Bayesian analysis. On the one hand, the frontoparietal structures (DLPFC, IPL, TPJ) are engaged not only in Go/NoGo tasks but also in experiments examining task switching, resolution of cognitive conflicts, working memory, and attention focusing (“multiple demand system” (Duncan, 2010); “task-general network” (Cole et al., 2014); “extrinsic mode network” (Hugdahl et al., 2015)). On the other hand, the most prominent feature of brain activity during Go/NoGo tasks compared to diverse non-inhibitory tasks is a right-dominant activity in DLPFC (meta-analyses: Buchsbaum et al., 2005, Nee, Wager & Jonides, 2007, Levy & Wagner, 2011, Cieslik et al., 2015, Zhang, Geng & Lee, 2017). Among different response inhibition tasks, several meta-analyses showed that an action-restraint (Go/NoGo task) elicits stronger activation in the right DLPFC and parietal cortex compared to another inhibition process, namely, action cancellation (stop-signal task) (Swick, Ashley & Turken, 2011, Cieslik et al., 2015, Zhang, Geng & Lee, 2017). Furthermore, “response uncertainty” tasks induced by the equal probability of Go and NoGo stimuli evoked more activity in the right DLPFC (near the cluster revealed in the current study with coordinates [x=45 y=33 z=28], see Table 2) compared to Go/NoGo and stop-signal tasks with a low probability of inhibitory stimuli (“response override” tasks) (Levy & Wagner, 2011).

In meta-analyses of Simmonds, Pekar & Mostofsky (2008) and Criaud & Boulinguez (2013), it was supposed that the increased activity in the right DLPFC observed during Go/NoGo tasks may be due to increased demands caused by working-memory and attentional load rather than the inhibition processes per se. The authors explained it by the fact that most of the fMRI studies used complex designs. Thus, it would be relatively more difficult to identify a NoGo stimulus, which is believed to increase the need for inhibitory control and consequently increase the brain inhibitory activity in NoGo trials. However, this confounding effect was controlled in the current fMRI study because we utilized a simple equiprobable Go/NoGo task.

Another prefrontal structure related to non-selective response inhibition in the current study was the left IFG (opercular part, AIFO). At the same time, the right IFG is commonly associated with response inhibition (Aron, Robbins & Poldrack, 2014). Based on the study of patients with brain damage (Aron et al., 2003) and other neuroimaging studies using primarily the stop-signal tasks (Aron, Robbins & Poldrack, 2014), it was claimed that the right IFG represents a key node of the response inhibition brain system. Currently, this opinion is under active discussion (Mostofsky & Simmonds, 2008, Hampshire et al., 2010, Hampshire, 2015a, Hampshire & Sharp, 2015b, Swick, Ashley & Turken, 2008, Swick & Chatham, 2014). It is noted that Aron et al. predominantly relied upon studies using the stop-signal task and did not consider studies using Go/NoGo tasks (Swick & Chatham, 2014). The studies by Swick, Ashley & Turken (2008) and Kramer et al. (2013) revealed worse performance in the Go/NoGo task when the left (not right) IFG was damaged. According to these findings, the left IFG can also be involved in inhibitory control during action restraint, which is additionally confirmed by our results.

Regarding the practically equivalent involvement of PMC in both Go and NoGo trials, it was previously demonstrated to participate in the planning and coordination of actions because its electrical stimulation results in involuntary motor actions (Desmurget et al., 2009). According to Duque et al. (2012), premotor cortex function is associated with inhibition of any premature actions and control of the time of action execution. The authors refer to this brain mechanism as “impulse control”, a concept that is similar to non-selective response inhibition (Criaud et al., 2017).

In summary, it can be noted that the selectivity of response inhibition has been more extensively studied to date than its non-selectivity. A considerable amount of research on inhibitory control has used tasks with a low probability of inhibitory stimuli. However, over time, it has become apparent that under several conditions, non-selectivity of response inhibition can, in principle, occur. Studies on non-selectivity of response inhibition typically focus on the possible brain mechanisms for non-selective (“global”) inhibition of motor responses in the settings of interference among multiple concurrent response options (De Jong, Coles & Logan, 1995, Coxon, Stinear & Byblow, 2007, Aron & Verbruggen, 2008, Frank, 2006, Duque & Ivry, 2009, Duque et al., 2010, MacDonald et al., 2017). Recently, the possibility of involvement of non-selective response inhibition not only for NoGo stimuli but also for Go stimuli was supposed by Criaud et al. (2017). In the current study, the practical equivalence of BOLD signals evoked by equiprobable NoGo and Go stimuli was demonstrated for a number of brain areas associated with response inhibition in an uncertain context, according to our meta-analysis. The present study results proved that response inhibition can act as a non-selective mechanism of action inhibition when the context is uncertain. Thus, one promising area of further research of brain mechanisms of response inhibition (and inhibitory control in general) is the study of the interplay between selective and non-selective inhibition as a function of the contextual uncertainty degree.

## 5. Conclusion

For the first time, combining a meta-analysis and second-level Bayesian analysis yielded results favouring the existence of non-selective response inhibition. In the present work, selectivity means that inhibition is triggered only by an inhibitory stimulus. The overlap between brain areas previously associated with response inhibition in uncertain context and brain areas demonstrating the practical equivalence of neuronal activity in equiprobable Go and NoGo trials was observed in the right DLPFC, IPL, TPJ, FEF, PMC, and left IFG (AIFO). When a subject was waiting for an equiprobable Go or NoGo stimulus, a non-selective inhibitory control process occurred in both Go trials and NoGo trials in opposition to the model of selective response inhibition. This type of response inhibition prevents the performance of any premature motor actions and operates in a non-selective, “global” mode. Its involvement is favoured by contextual uncertainty caused by the equally probable presentation of Go and NoGo stimuli. Presumably, upon the identification of a Go stimulus, the inhibition is released, and the process of action execution is initiated, i.e., it acts as a gating mechanism for accessing a prepared motor program. At the same time, we do not discard the opportunity of involvement of selective inhibitory mechanisms in a less uncertain context. Therefore, future research should address the issue of how brain mechanisms of selective and non-selective response inhibition are inter-related.

## 6. Limitations and further considerations

It is important to note that we looked for practically equivalent increases in regional brain activity as measured by BOLD-signal. Firstly, it is possible to look for the similarity of local activity patterns by employing multivariate classification techniques in future studies. Secondly, due to the limited temporal resolution of BOLD fMRI, it is necessary to combine it with electrophysiological methods. In particular, evidence for the practical equivalence of early ERP amplitudes (170 ms) revealed previously in equiprobable NoGo and Go trials should be provided. Thirdly, future research should consider how non-selective response inhibition is mediated by interactions between the revealed cortical structures and subcortical structures that form inhibitory and excitatory cortico-striato-thalamo cortical circuits.

## 7. Data and code availability statements

Matlab codes used to convert beta-values to PSC and perform Bayesian inference are available in Github (https://github.com/Masharipov/Non-selective-inhibition). Meta-analysis Z-score map and thresholded ALE map, Bayesian analysis and conjunction analysis results are available in Neurovault (https://neurovault.org/collections/6009/). De-identified neuroimaging data, including all individual contrasts, are available upon reasonable request by message to the corresponding author. Usage and sharing of these data must comply with ethics approval of the N.P. Bechtereva Institute of the Human brain of the Russian Academy of Sciences.

## 8. Acknowledgments

Current study in a part for Bayesian analysis of fMRI data was supported by the Russian Science Foundation grant #19-18-00454. The meta-analysis of Go/NoGo fMRI studies was performed within the state assignment of the Ministry of Education and Science of Russian Federation (theme number AAAA-A19-119101890066-2).

