## Supplementary materials for "Non-selective response inhibition in equiprobable Go/NoGo task: Bayesian analysis of fMRI data"

Phone number: +7 812 670-09-51

Fax number: +7 812 2343247

### **Table S1. The characteristics of studies included in the meta-analysis**

| Study (Author) | Sample (M/F) | Field  (Tesla) | Age  (mean) | Design | Contrast | Foci | Space |
| --- | --- | --- | --- | --- | --- | --- | --- |
| Menon et al. (2001) | 8/6 | 1.5 | 17-41  (23.6) | 12 alternating 26 s blocks of  Go/NoGo conditions  (50% “X” letter – NoGo stimuli)  and Go-control (no “X” letter) | Go/NoGo > Go-control blocks | 13 | MNI (SPM97) |
| Booth et al. (2003) | 5/7 | 1.5 | 20-30  (25.1) | 12 alternating 36 s blocks of  Go/NoGo conditions  (50% Red triangle – NoGo stimuli)  and Go-control (all stimuli – Go) | Go/NoGo > Go-control blocks  (within adult group) | 13 | MNI (SPM99) |
| Horn et al. (2003) | 19 M | 1.5 | 18-50* | 6 alternating 37.9 s blocks of  Go/NoGo conditions (50% “V” letter – NoGo stimuli)  and Go-control (no “V” letter) | Go/NoGo > Go-control blocks | 14 | MNI (SPM99) |
| Maguire et al. (2003) | 6 M | 1.5 | 22-30* | 20 alternating 24 s blocks of  Go/NoGo conditions (50% Red square – NoGo stimuli)  and Go-control (no Red square) | Go/NoGo > Go-control blocks | 6 | MNI (SPM99) |
| Asahi et al. (2004) | 10/7 | 1.5 | 23-30  (25.1) | 8 alternating 36 s blocks of  Go/NoGo conditions (50% “X” letter – NoGo stimuli)  and Go-control (no “X” letter) | Go/NoGo > Go-control blocks | 11 | MNI (SPM99) |
| Völlm et al. (2004) | 8 M | 1.5 | (29.9)* | 8 alternating 44.2 s blocks of  Go/NoGo conditions (50% “V” letter – NoGo stimuli)  and Go-control (no “V” letter) | Go/NoGo > Go-control blocks  (within healthy control) | 13 | MNI (SPM2) |
| Altshuler et al. (2005) | 5/8 | 3 | (31)* | 8 alternating 30.5 s blocks of  Go/NoGo conditions (50% “X” letter – NoGo stimuli)  and Go-control (all stimuli – Go) | Go/NoGo > Go-control blocks  (within healthy control) | 4 | TAL (AIR) |
| Den-Ben et al. (2005) | 12 M | 1.5 | 19-36  (24.7) | 8 alternating 45 s blocks of  Go/NoGo conditions (50% “V” letter – NoGo stimuli)  and Go-control (no “V” letter);  placebo and citalopram sessions | Go/NoGo > Go-control blocks  (main effect of task) | 15 | MNI (SPM2) |
| Brown et al. (2006) | 21/37 | 3 | (45.3)* | 6 alternating 30 s blocks of  Go/NoGo conditions (50% “V” letter – NoGo stimuli)  and Go-control (no “V” letter) | Go/NoGo > Go-control blocks | 5 | MNI (SPM2) |
| Passamonti et al. (2006) | 24 M | 1.5 | (30.25) | 4 alternating 28 s blocks of  Go/NoGo conditions (50% “V” letter – NoGo stimuli)  and Go-control (no “V” letter) | Go/NoGo > Go-control blocks  (independently of genotype) | 9 | MNI (SPM99) |
| Völlm et al. (2006) | 45 M | 1.5 | 18-45  (27.3) | 8 alternating 44.2 s blocks of  Go/NoGo conditions (50% “V” letter – NoGo stimuli)  and Go-control (no “V” letter) | Go/NoGo > Go-control blocks | 17 | MNI (SPM2) |
| Mobbs et al. (2007) | 2/9 | 1.5 | 19-54  (30.2) | 12 alternating 26 s blocks of  Go/NoGo conditions (50% “X” letter – NoGo stimuli)  and Go-control (no “X” letter) | Go/NoGo > Go-control blocks  (within healthy control) | 4 | MNI (SPM99) |
| Dillo et al. (2010) | 11/4 | 1.5 | 21-46  (28.8) | 10 alternating 21 s blocks of  Go/NoGo conditions (50% “X” letter – NoGo stimuli)  and Go-control (all stimuli – Go) | Go/NoGo > Go-control blocks  (within healthy control) | 2 | MNI (SPM2) |
| Stokes et al. (2011) | 22/27 | 3 | (37)* | 10 alternating 36 s blocks of  Go/NoGo conditions (~50% “V” letter – NoGo stimuli)  and Go-control (no “V” letter) | Go/NoGo > Go-control blocks  (independently of genotype) | 6 | TAL (SPM8, Brett’s nonlinear transf.) |
| Townsend et al. (2012) | 17/13 | 3 | (37)* | 8 alternating 30.5 s blocks of  Go/NoGo conditions (50% “X” letter – NoGo stimuli)  and Go-control (no “X” letter) | Go/NoGo > Go-control blocks  (within healthy control) | 24 | MNI (FSL 4.0) |
| Chen et al. (2014) | 15 M | 3 | (24.5)* | 12 alternating 30 s blocks of  Go/NoGo conditions (50% “0” number – NoGo stimuli)  and Go-control (no “0” number) | Go/NoGo > Go-control blocks  (within healthy control) | 7 | MNI (SPM5) |
| Liu et al. (2014) | 11 M | 3 | (22.5)* | 8 alternating 30 s blocks of  Go/NoGo conditions (50% “pentagons” – NoGo stimuli)  and Go-control (no “pentagons”) | Go/NoGo > Go-control blocks  (within healthy control) | 2 | TAL (SPM5, linear transf.) |
| Penfold et al. (2015) | 10/10 | 3 | (35.6)* | 8 alternating 30 s blocks of  Go/NoGo conditions (50% “X” letter – NoGo stimuli)  and Go-control (no “X” letter) | Go/NoGo > Go-control blocks  (within healthy control) | 30 | MNI (FSL 5.0.4) |
| Shafritz et al. (2015) | 12/3 | 3 | 12-23  (18.4) | 8 alternating 30 s blocks of  Go/NoGo conditions (50% “X” letter – NoGo stimuli)  and Go-control (no “X” letter) | Go/NoGo > Go-control blocks  (within healthy control; non-emotional task) | 5 | MNI (SPM8) |
| Pornpattananangkul et al. (2016) | 29/29 | 3 | (23)* | 8 alternating 30 s blocks of  Go/NoGo conditions (Caucasian and Japanese Americans: 50% “V” letter – NoGo stimuli; Native Japanese: 50% “レ” letter – NoGo stimuli)  and Go-control (no “V”/“レ” letter) | Go/NoGo > Go-control blocks  (across Caucasian  Americans, Japanese Americans, and Native Japanese) | 10 | MNI (SPM8) |

* - mean age or age range not reported.

MNI - Montreal Neurological Institute space; TAL – Talairach space; SPM - spatial normalization were performed with Statistical Parametric Mapping tools; FSL - spatial normalization were performed with Functional Magnetic Resonance Imaging of the Brain (FMRIB) Software Library; AIR - spatial normalization were performed with Automated Image Registration tools.

### **Power analysis**

The effect sizes typical of selective inhibition were estimated using UCLA Consortium for Neuropsychiatric Phenomic dataset (openneuro.org/datasets/ds000030) (Poldrack et al., 2016). A total of 115 subjects were included in the analysis (55 females, 60 males, mean age = 31.7 ± 8.9 years) after removing subjects with no data for the stop-signal task, high level (>15%) of errors in the Go-trials and reported to have problems with the raw data (Gorgolewski et al., 2017). The minimal preprocessing pipeline for the UCLA dataset described in (Gorgolewski et al., 2017). Spatial smoothing was applied with 8 mm FWHM (full width at half maximum) Gaussian smoothing kernel to the preprocessed images. Spatial smoothing performed using SPM12. At the first level of analysis, the frequentist parameter estimation was applied to calculate the beta values of the general linear models using SPM12. The stop-signal task consisted of 96 Go trials and 32 Stop-signal trials (approximately half of them were correct). Jittered null events separated every trial. Three regressors were included in a first-level model with fixed event duration of 1.5 s. Twenty-four head motion regressors were included in each first-level model (6 head motion parameters, 6 head motion parameters one-time point before, and 12 corresponding squared items) to minimize the head motion artefacts (Friston et al., 1996). We performed a one-sample test on the linear contrasts “Correct-Stop > Go” created at the first level of analysis (see Fig.S1).

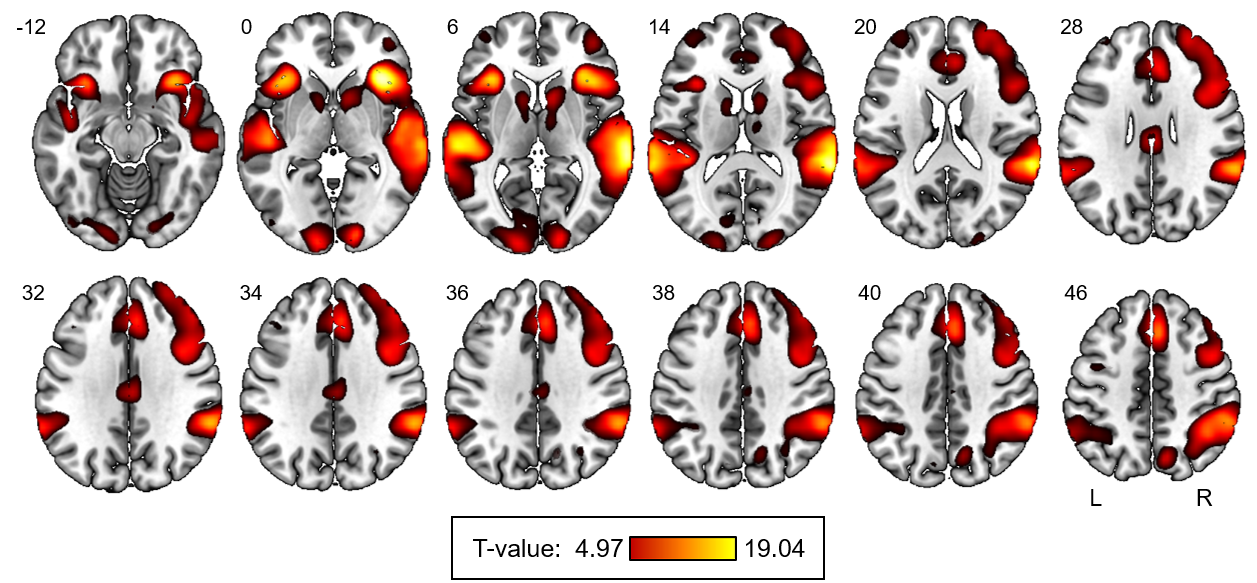

### **Figure S1. The result for the selective inhibition contrast (“Correct-Stop > Go” effect)**

Classical inference, with an FWE-corrected voxel-wise threshold of p < 0.05, was used. Cohen’s d were calculated from the T-values. The mean effect sizes were estimated within the cortical brain regions revealed by the current meta-analysis: (1) right dorsolateral prefrontal cortex (d = 0.70), (2) right inferior parietal lobule (d = 0.57), (3) right temporoparietal junction (d = 1.12), (4) right inferior frontal gyrus and anterior insula (d = 1.05), (5) left inferior frontal gyrus and anterior insula (d = 1.01), (6) bilateral anterior cingulate cortex and supplementary motor area (d = 0.77). See the power curve for the minimum estimated effect size (d = 0.57) on Fig. S2.

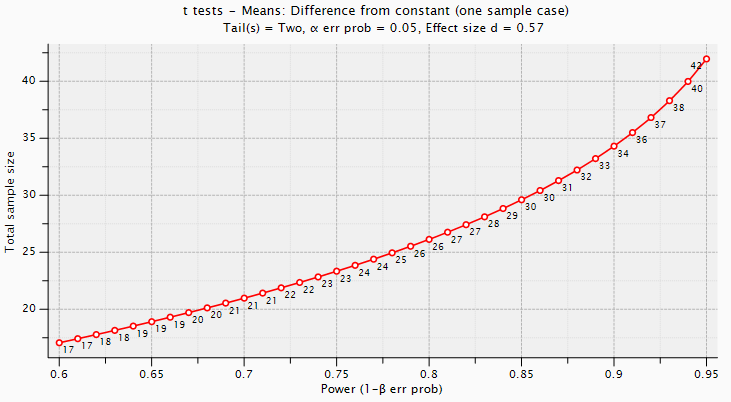

### **Figure S2. The power curve for d = 0.57, two-tailed one-sample test, alpha = 0.05**

The plot was obtained using GPower 3.1.9.7.

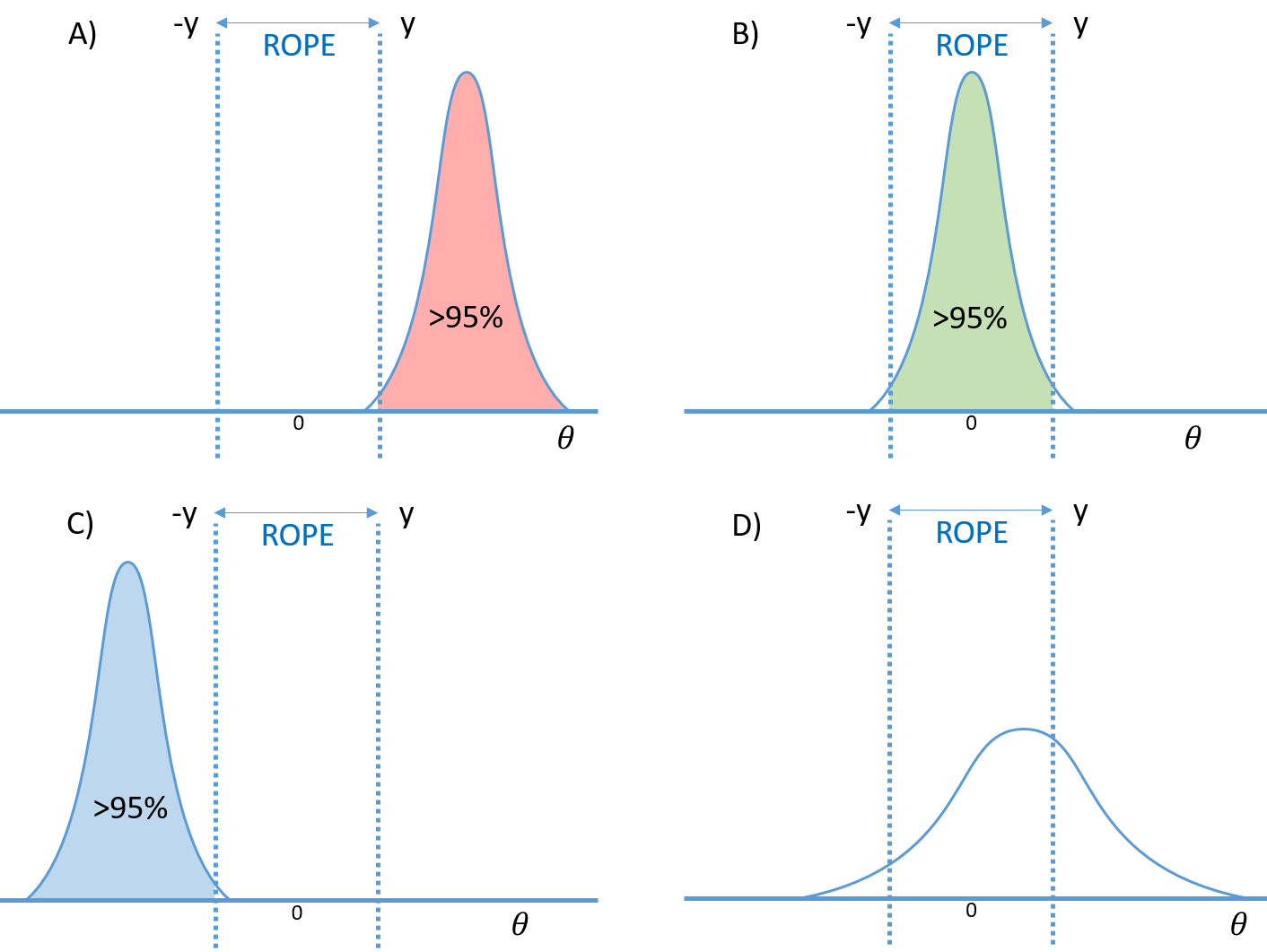

### **Figure S3. Bayesian parameter inference with the “ROPE-only” decision rule**

Four possible variants of the posterior distribution of the effect (*θ = cβ*) are presented. A) If the posterior probability of the effect exceeding the effect size threshold, *γ*, is greater than the predefined probability threshold, *α* = 95%, then the hypothesis on the presence of the positive effect can be accepted. B) If the effect value falls within the interval [-*γ*; *γ*] or the region of practical equivalence (ROPE) with a probability of *α* = 0.95, then the null hypothesis can be accepted. C) If the posterior probability of the effect less than -*γ* is greater than 95%, then the hypothesis on the presence of the negative effect can be accepted. D) The posterior probability of detecting an effect inside or outside the ROPE does not exceed 95%. The data in the particular voxel are insufficient to distinguish the null hypothesis from the alternative hypothesis (low-confidence).

### **Table S2. The result of classical voxel-wise inference for the “Go > NoGo” effect**

Voxel-wise pFWE<0.05.

| № | Cluster size, mm^3^ | Peak MNI-coordinate, mm | Peak t-value | Anatomical localization (L – left, R – right hemisphere; BA – Brodmann area) |
| --- | --- | --- | --- | --- |
| 1 | 257688 | -39 -1 -1 | 17.55 | L: Insula, IFG (AIFO), BA 13, 44, 47 |
|  |  | -48 -25 44 | 16.63 | L: Postcentral, IPL, BA 2, 3, 4, 40 |
|  |  | 57 11 17 | 16.26 | R: IFG, Precentral, BA 9, 44 |
|  |  | -57 -22 17 | 16.01 | L: Postcentral |
|  |  | 9 -55 -16 | 15.87 | L/R: Vermis, R: Cerebellum lobe |
|  |  | 39 5 -1 | 15.76 | R: Insula, BA 13 |
|  |  | -6 -25 41 | 14.18 | L/R: MCC, BA 24 |
|  |  | -3 5 38 | 14.02 | L/R: MCC, SMA, BA 6, 24 |
|  |  | -36 -19 56 | 14.01 | L: Precentral, Postcentral, BA 2, 3, 4, 9 |
|  |  | -15 -22 8 | 13.70 | L: Thalamus |
|  |  | -21 -43 71 | 13.03 | L: SPL, BA 5, 7 |
|  |  | 12 2 8 | 12.71 | R: Caudate, putamen, pallidum |
|  |  | 3 -1 53 | 12.63 | L/R: SMA, BA 6 |
|  |  | 51 11 5 | 12.12 | R: IFG (AIFO), BA 13, 44, 45, 47 |
|  |  | 3 -31 -1 | 9.62 | L/R: Thalamus |
|  |  | -54 -43 32 | 9.29 | L: TPJ (supramarginal gyrus), BA 40 |
|  |  | -9 11 -1 | 9.13 | L: Caudate, putamen, pallidum |
|  |  | -54 8 23 | 8.83 | L: IFG, Precentral, BA 9, 44 |
|  |  | 42 41 5 | 8.78 | R: IFG, BA 46 |
|  |  | 15 5 65 | 8.71 | R: SFG, BA 6 |
|  |  | 39 44 8 | 8.71 | R: MFG, BA 10 |
|  |  | 3 38 32 | 7.31 | L/R: ACC, BA 32 |
|  |  | 33 20 -13 | 7.93 | R: AIFO, BA 13, 44, 45, 47 |
|  |  | 24 -40 68 | 6.13 | R: Postcentral, BA 2, 3, 5 |
| 2 | 26055 | 57 -34 41 | 14.33 | R: TPJ (supramarginal gyrus), BA 40 |
|  |  | 60 -19 26 | 11.89 | R: Postcentral, BA 2, 3 |
|  |  | 42 -40 56 | 8.35 | R: IPL, BA 40 |
| 3 | 2403 | 21 56 26 | 7.87 | R: SFG, MFG, BA 10 |
| 4 | 1998 | -51 -64 2 | 7.00 | L: Middle temporal gyrus, BA 22, 37 |
| 5 | 1512 | -36 38 29 | 8.11 | L: MFG, BA 10 |
| 6 | 594 | 18 -49 68 | 6.24 | R: Postcentral, BA 2, 5 |
| 7 | 513 | 57 -55 2 | 6.29 | R: Middle temporal gyrus, BA 22, 37 |

Abbreviations: SFG – superior frontal gyrus, MFG – middle frontal gyrus, IFG – inferior frontal gyrus, AIFO – anterior insula/frontal operculum, TPJ – temporoparietal junction, SMA – supplementary motor area, ACC – anterior cingulate cortex, MCC – middle cingulate cortex, IPL – inferior parietal lobule, SPL – superior parietal lobule.

### **Table S3. The result of Bayesian parameter inference for the “Go > NoGo” effect**

*γ* = 1 Prior SD = 0.1%, LogOdds > 3 (*PP* > 0.95).

| № | Cluster size, mm^3^ | Peak MNI-coordinate, mm | Peak LogOdds | Anatomical localization (L – left, R – right hemisphere; BA – Brodmann area) |
| --- | --- | --- | --- | --- |
| 1 | 161892 | -36 -19 56 | 36.04 | L: Precentral, Postcentral, BA 2, 3, 4, 9 |
|  |  | 3 -1 53 | 36.04 | L/R: SMA, BA 6 |
|  |  | -3 -25 44 | 36.04 | L/R: MCC, BA 24 |
|  |  | -48 -25 44 | 36.04 | L: Postcentral, IPL, BA 2, 3, 4, 40 |
|  |  | -3 5 38 | 36.04 | L/R: MCC, SMA, BA 6, 24 |
|  |  | -51 -25 20 | 36.04 | L: Postcentral, |
|  |  | -57 -22 17 | 36.04 | L: Postcentral, TPJ (supramarginal), BA 40 |
|  |  | -39 -1 -1 | 36.04 | L: Insula, IFG (AIFO), BA 13, 44, 47 |
|  |  | 6 -55 -10 | 36.04 | L/R: Vermis |
|  |  | 18 -52 -22 | 36.04 | R: Cerebellum lobe |
|  |  | -12 -22 8 | 35.64 | L: Thalamus |
|  |  | -24 -46 71 | 33.96 | L: SPL, BA 5, 7 |
|  |  | 12 2 8 | 26.26 | R: Caudate, putamen, pallidum |
|  |  | 3 -31 -1 | 18.86 | L/R: Thalamus |
|  |  | -54 8 23 | 14.42 | L: IFG, Precentral, BA 9, 44 |
|  |  | -9 8 2 | 12.53 | L: Caudate, putamen, pallidum |
|  |  | 3 38 29 | 6.30 | L/R: ACC, BA 32 |
| 2 | 20925 | 60 -19 26 | 28.78 | R: Postcentral, BA 2, 3 |
|  |  | 54 -25 50 | 13.09 | R: Postcentral, BA 2, 3 |
|  |  | 42 -40 56 | 12.56 | R: IPL, TPJ (supramarginal gyrus), BA 40 |
|  |  | 27 -37 71 | 3.90 | R: Postcentral, BA 2 |
| 3 | 17010 | 42 5 -4 | 36.04 | R: Insula, BA 13 |
|  |  | 57 11 14 | 36.04 | R: IFG (AIFO), Precentral, BA 9, 44, 45, 47 |
| 4 | 1755 | 42 47 8 | 8.29 | R: MFG, IFG, BA 10, 46 |
| 5 | 351 | -36 41 26 | 4.73 | L: MFG, BA 10 |
| 6 | 270 | 24 56 26 | 4.92 | R: SFG, MFG, BA 10 |

Abbreviations: SFG – superior frontal gyrus, MFG – middle frontal gyrus, IFG – inferior frontal gyrus, AIFO – anterior insula/frontal operculum, TPJ – temporoparietal junction, SMA – supplementary motor area, ACC – anterior cingulate cortex, MCC – middle cingulate cortex, IPL – inferior parietal lobule, SPL – superior parietal lobule.

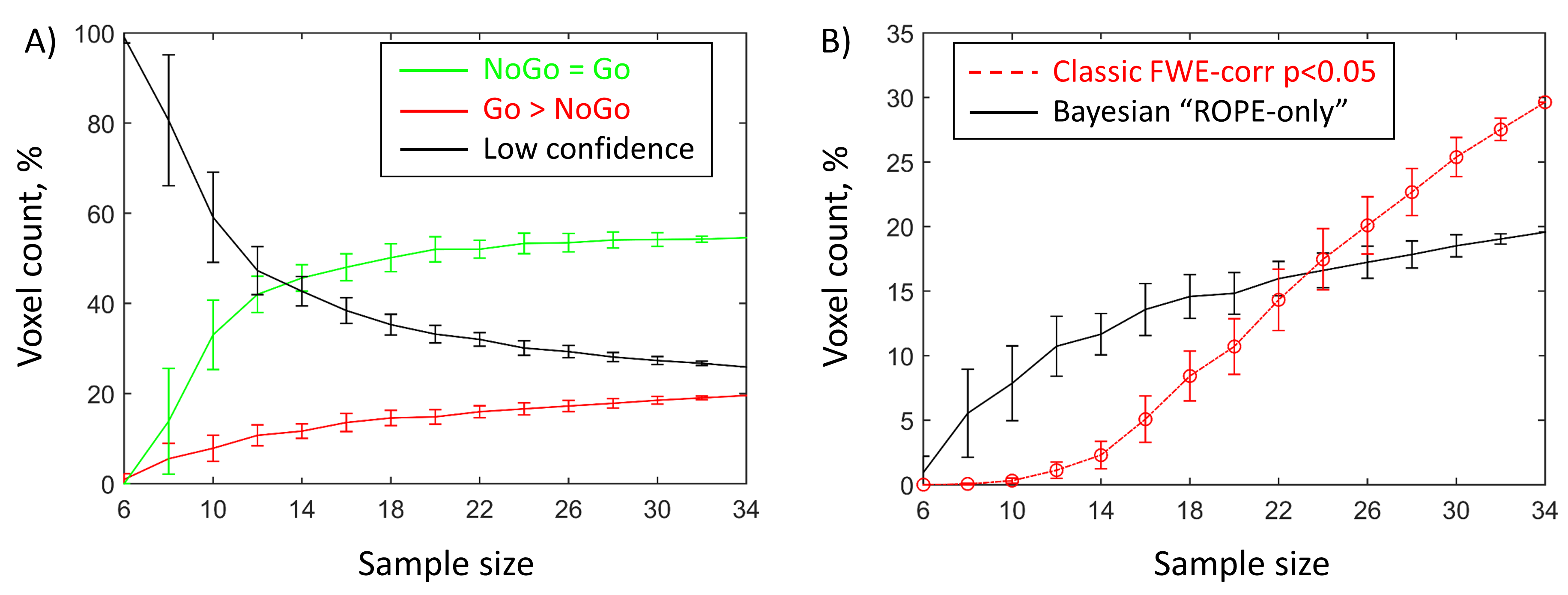

### **Figure S4. Sample size dependencies for Bayesian and classical inference**

To estimate how the sample size influences the results, we performed Bayesian and NHST-based analysis for the samples of different sizes: from 6 to 34 subjects with a step of 2 subjects. Thirty random groups were sampled for each step. Error bars represent mean and standard deviation across thirty random groups.

A) Bayesian parameter inference results. Dependencies between sample size and the number of “NoGo = Go”, “Go > NoGo”, and “low confidence” voxels (in % from the total number of voxels) are depicted by the green, red and black colours, respectively. An enlargement of the sample size led to an increase in the number of “NoGo = Go” and “Go > NoGo” voxels, and a decrease in the number of ‘low confidence’ voxels. The largest gain in the number of “NoGo = Go” and “Go > NoGo” voxels can be noted from 6 to 20 subjects. After 20 subjects, all dependencies reach a plateau.

B) The number of “Go > NoGo” voxels revealed using classical inference with pFWE<0.05 (red colour) and Bayesian inference (black colour). Bayesian parameter inference based on the “ROPE-only” decision rule is more sensitive to “activations” than classical inference with the voxel-wise FWE correction for relatively small sample sizes. The number of “Go > NoGo” voxels revealed using classical NHST with FWE-correction showed a steady linear increase with increasing sample size (see Fig. S4B). At larger sample sizes, there will be a discrepancy between classical and Bayesian inference, which is a manifestation of the Jeffreys-Lindley paradox (Jeffreys, 1939/1948; Lindley, 1957; Friston, 2012). These dependencies illustrate that Bayesian parameter inference protects from detecting “trivial” effects that can be appeared as a result of increased sample size if the classical (frequentist) inference is used (Friston et al., 2012).

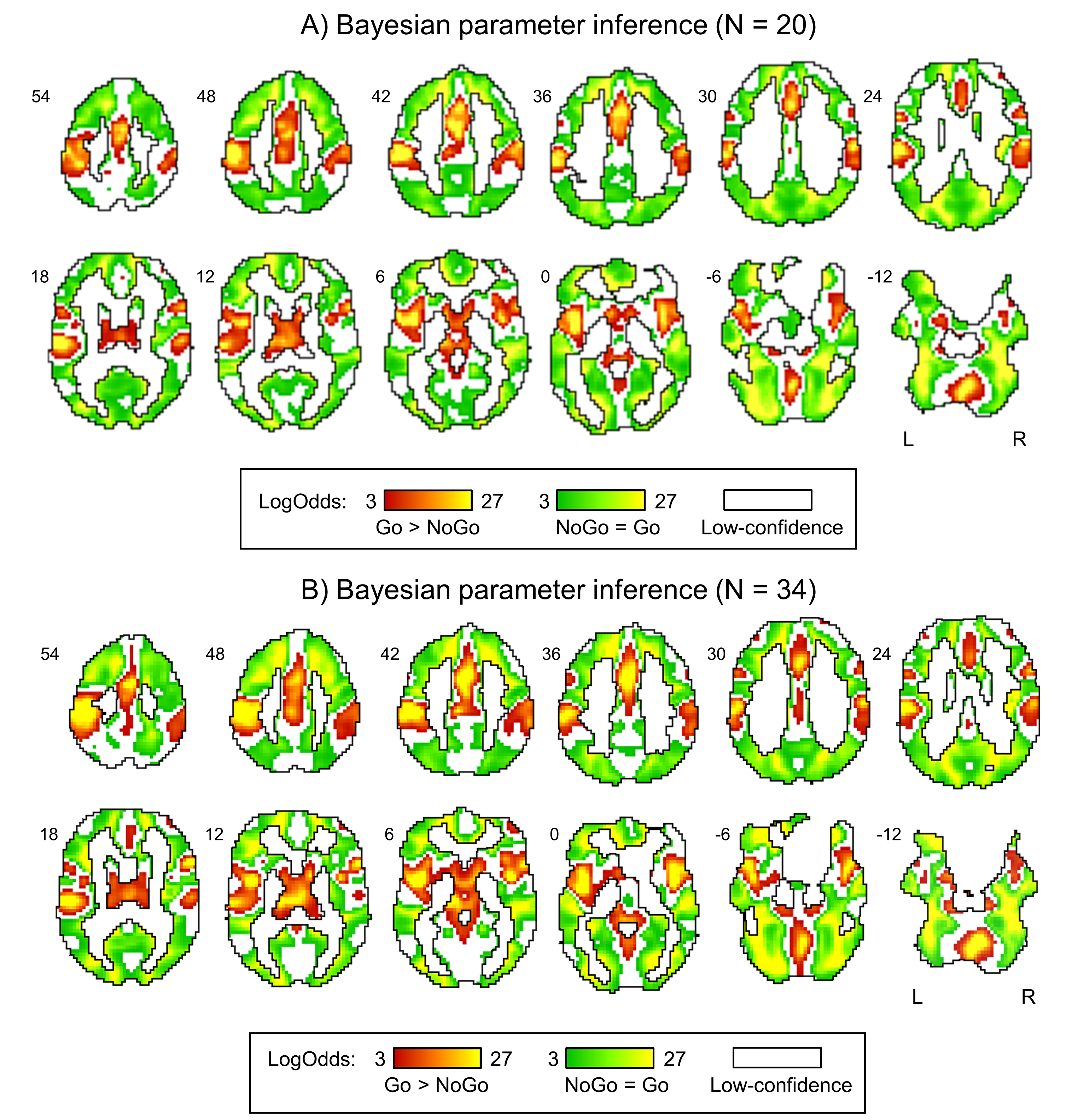

### **Figure S5. The result of Bayesian parameter inference for the “NoGo vs. Go” comparison obtained using the sample size of 20 and 34 subjects**

A) Results for the sample size N = 20. B) Results for the sample size N = 34.

The results of Bayesian parameter inference obtained using the sample size of 20 subjects were similar to those obtained using 34 subjects. Dice coefficients for the thresholded “NoGo = Go” and “Go < NoGo” images obtained using 20 and 34 subjects were 0.88 and 0.85, respectively. The effect size threshold *γ* = 1 prior SD*_θ_* = 0.1% and LogOdds > 3 (*PP* > 0.95). The “NoGo > Go” effect was not revealed. Red colour depicts the “Go > NoGo” effect. Green colour depicts the “NoGo = Go” effect. White colour depicts “low-confidence” voxels.

### **Table S4. The result of Bayesian parameter inference for the “Go > NoGo” effect using the sample size of 20 subjects**

*γ* = 1 Prior SD = 0.1%, LogOdds > 3 (*PP* > 0.95).

| № | Cluster size, mm^3^ | Peak MNI-coordinate, mm | Peak  LogOdds | Anatomical localization (L – left, R – right hemisphere; BA – Brodmann area) |
| --- | --- | --- | --- | --- |
| 1 | 34425 | 3 -61 -13 | 36.04 | L/R: Vermis |
|  |  | 18 -61 -19 | 35.64 | R: Cerebellum lobe |
|  |  | -12 -19 8 | 29.81 | L: Thalamus |
|  |  | 12 2 8 | 21.39 | R: Caudate, putamen, pallidum |
|  |  | -9 2 5 | 21.32 | L: Caudate, putamen, pallidum |
|  |  | 6 -16 11 | 16.50 | R: Thalamus |
| 2 | 93879 | -48 11 -7 | 36.04 | L: AIFO, BA 13, 44, 47 |
|  |  | -39 -1 -4 | 36.04 | L: Insula, BA 13 |
|  |  | -48 -28 41 | 36.04 | L: Postcentral, BA 2, 3 |
|  |  | -54 -22 20 | 36.04 | L: Postcentral, BA 2, 3 |
|  |  | 0 14 38 | 36.04 | L/R: ACC, BA 32 |
|  |  | 0 17 26 | 33.85 | L/R: ACC, BA 24 |
|  |  | 3 -10 41 | 32.46 | L/R: MCC, BA 24 |
|  |  | -21 -43 71 | 31.74 | L: SPL, BA 5, 7 |
|  |  | -36 -34 59 | 30.70 | L: Postcentral, BA 2, 3, 4 |
|  |  | 3 -4 56 | 27.98 | L/R: SMA, BA 6 |
|  |  | -54 8 20 | 26.19 | L: IFG, BA 44 |
| 3 | 14310 | 39 8 -1 | 35.64 | R: AIFO, BA 13, 47 |
|  |  | 57 11 14 | 31.24 | R: IFG, BA 44 |
|  |  | 42 -1 5 | 25.81 | R: Insula, BA 13 |
| 4 | 19278 | 57 -16 26 | 34.66 | R: Postcentral, BA 2, 3 |
|  |  | 57 -16 41 | 25.91 | R: Postcentral, BA 3 |
|  |  | 39 -37 50 | 15.89 | R: IPL, TPJ (supramarginal), BA 40 |
|  |  | 57 -34 26 | 14.13 | R: TPJ (supramarginal gyrus), BA 40 |
| 5 | 837 | 27 -7 65 | 6.43 | R: SFG, BA 6 |
| 6 | 270 | 36 47 23 | 4.36 | R: MFG, BA 10 |

Abbreviations: SFG – superior frontal gyrus, MFG – middle frontal gyrus, IFG – inferior frontal gyrus, AIFO – anterior insula/frontal operculum, TPJ – temporoparietal junction, SMA – supplementary motor area, ACC – anterior cingulate cortex, MCC – middle cingulate cortex, IPL – inferior parietal lobule, SPL – superior parietal lobule.

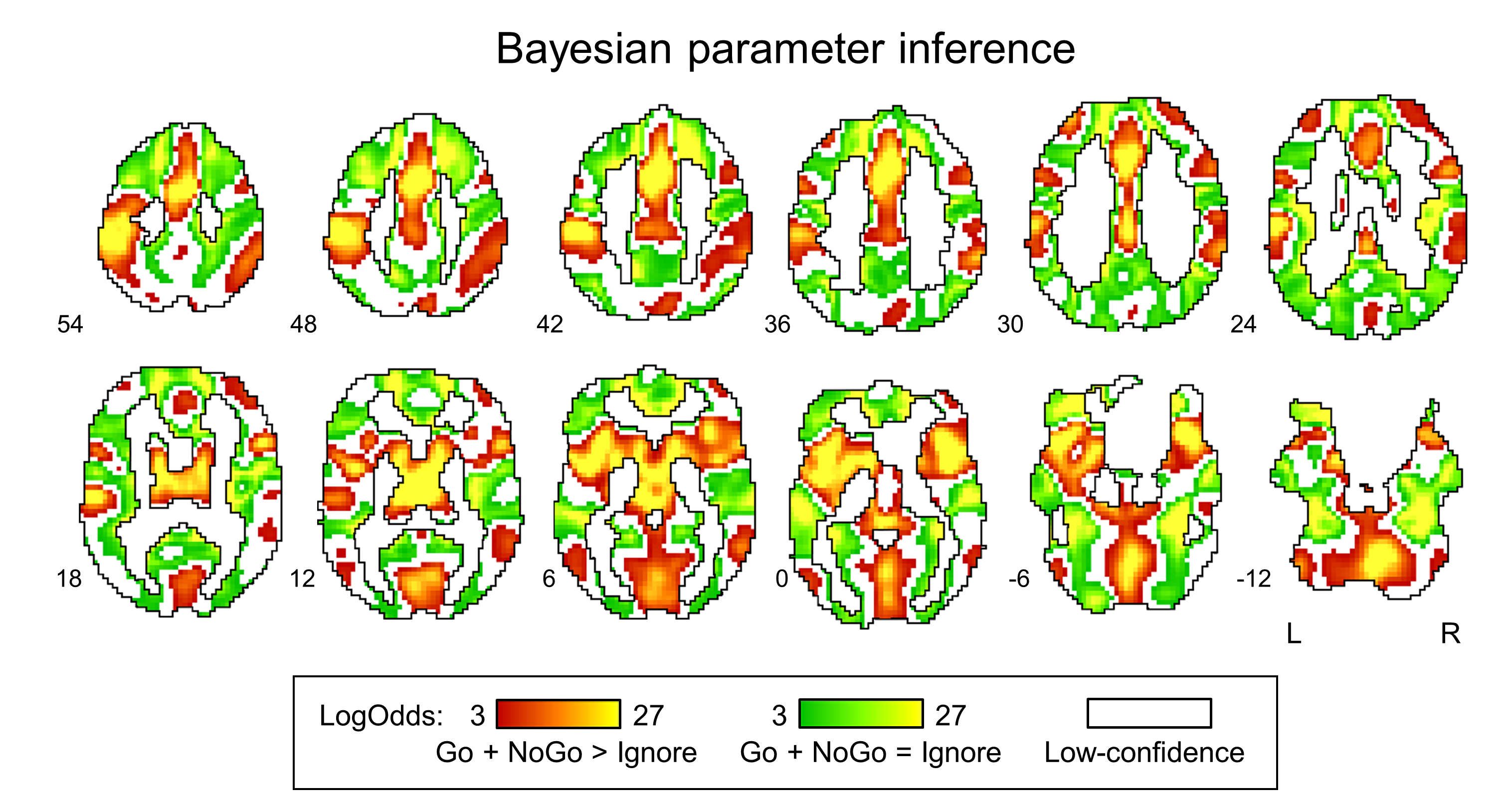

### **Figure S6. The result of Bayesian parameter inference for the “Go + NoGo > Ignore” comparison**

The effect size threshold *γ* = 1 prior SD*_θ_* = 0.9% and LogOdds > 3 (*PP* > 0.95). The “Ignore > Go + NoGo” effect was not revealed. Red colour depicts the “Go + NoGo > Ignore” effect. Green colour depicts the “Go + NoGo = Ignore” effect. White colour depicts “low-confidence” voxels.

### **Table S5. The result of Bayesian parameter inference for the “Go + NoGo > Ignore” effect**

*γ* = 1 Prior SD = 0.1%, LogOdds > 3 (*PP* > 0.95).

| № | Cluster size, mm^3^ | Peak MNI-coordinate, mm | Peak  LogOdds | Anatomical localization (L – left, R – right hemisphere; BA – Brodmann area) |
| --- | --- | --- | --- | --- |
| 1 | 273159 | -48 -28 50 | 36.04 | L: Precentral, Postcentral, BA 2, 3, 4 |
|  |  | 0 2 44 | 36.04 | L/R: MCC, SMA, BA 6, 24, 32 |
|  |  | 6 23 29 | 36.04 | L/R: ACC, MCC, BA 24 |
|  |  | -39 17 -4 | 36.04 | L: Insula, IFG (AIFO), BA 13, 44, 47 |
|  |  | 24 -52 -22 | 36.04 | R: Cerebellum lobe |
|  |  | 39 14 -1 | 36.04 | R: Insula, IFG (AIFO), BA 13, 44, 47 |
|  |  | -42 -4 2 | 36.04 | L: Insula, BA 13 |
|  |  | -57 -25 20 | 36.04 | L: Postcentral, TPJ (supramarginal), BA 40 |
|  |  | -9 -19 8 | 36.04 | L: Thalamus |
|  |  | 15 8 5 | 36.04 | R: Putamen, caudate, pallidum |
|  |  | 6 -19 8 | 36.04 | R: Thalamus |
|  |  | -12 2 14 | 36.04 | L: Caudate, putamen, pallidum |
|  |  | 39 5 -1 | 36.04 | R: Insula, BA13 |
|  |  | 3 -64 -7 | 36.04 | L/R: Vermis |
|  |  | 0 -25 29 | 31.32 | L/R: MCC, BA 23 |
|  |  | 9 -1 68 | 27.51 | L/R: SMA, BA 6 |
|  |  | -3 -79 11 | 24.71 | L/R: Cuneus, Lingual gyrus, BA 17, 18 |
|  |  | -18 -4 68 | 16.98 | L: SFG (FEF/PMC), Precentral, BA 6, 8 |
|  |  | 9 -73 47 | 11.74 | L/R: Precuneus, BA 7 |
|  |  | 39 -1 59 | 10.38 | R: SFG (FEF/PMC), Precentral, BA 6, 8 |
|  |  | -57 -40 29 | 9.46 | L: TPJ (supramarginal), BA 40 |
|  |  | 33 47 29 | 8.84 | R: SFG (DLPFC), BA 9 |
|  |  | 51 29 26 | 4.35 | R: MFG (DLPFC), BA 9 |
| 2 | 26676 | 60 -40 32 | 25.61 | R: TPJ (supramarginal), BA 40 |
|  |  | 42 -46 56 | 16.40 | R: SPL, IPL, BA 40 |
|  |  | 54 -52 5 | 15.55 | R: TPJ, STG, MTG, BA 22, 39 |
|  |  | 60 -19 29 | 12.20 | R: Postcentral, BA 3, 4 |
|  |  | 30 -64 44 | 5.11 | R: SPL, IPL, Angular, BA 7, BA 40 |
| 3 | 3024 | -36 44 29 | 20.64 | L: SFG, MFG (DLPFC), BA 10 |
| 4 | 513 | 51 -31 -4 | 4.77 | R: MTG, BA 21 |
| 5 | 162 | 33 -16 -4 | 3.44 | R: Putamen |

Abbreviations: SFG – superior frontal gyrus, MFG – middle frontal gyrus, IFG – inferior frontal gyrus, FEF – frontal eye field, PMC – premotor cortex, AIFO – anterior insula/frontal operculum, TPJ – temporoparietal junction, SMA – supplementary motor area, ACC – anterior cingulate cortex, MCC – middle cingulate cortex, IPL – inferior parietal lobule, SPL – superior parietal lobule, STG – superior temporal gyrus, MTG – middle temporal gyrus.

### **References (Studies included in the meta-analysis)**

Altshuler, L. L., Bookheimer, S. Y., Townsend, J., Proenza, M. A., Eisenberger, N., Sabb, F., … Cohen, M. S. (2005). Blunted Activation in Orbitofrontal Cortex During Mania: A Functional Magnetic Resonance Imaging Study. *Biological Psychiatry*, *58*(10), 763–769. https://doi.org/10.1016/j.biopsych.2005.09.012

Asahi, Sh., Okamoto, Y., Okada, G., Yamawaki, Sh., & Yokota, N. (2004). Negative correlation between right prefrontal activity during response inhibition and impulsiveness: A fMRI study. *European Archives of Psychiatry and Clinical Neurosciences*, *254*(4). https://doi.org/10.1007/s00406-004-0488-z

Booth, J. R., Burman, D. D., Meyer, J. R., Lei, Z., Trommer, B. L., Davenport, N. D., … Mesulam, M. M. (2003). Neural development of selective attention and response inhibition. *NeuroImage*, *20*(2), 737–751. https://doi.org/10.1016/S1053-8119(03)00404-X

Brown, S. M., Manuck, S. B., Flory, J. D., & Hariri, A. R. (2006). Neural basis of individual differences in impulsivity: Contributions of corticolimbic circuits for behavioral arousal and control. *Emotion*, *6*(2), 239–245. https://doi.org/10.1037/1528-3542.6.2.239

Chen, C.-Y., Huang, M.-F., Yen, J.-Y., Chen, C.-S., Liu, G.-C., Yen, C.-F., & Ko, C.-H. (2014). Brain correlates of response inhibition in Internet gaming disorder. *Psychiatry and Clinical Neurosciences*, *69*(4), 201–209. https://doi.org/10.1111/pcn.12224

Del-Ben, C. M., Deakin, J. F. W., Mckie, S., Delvai, N. A., Williams, S. R., Elliott, R., … Anderson, I. M. (2005). The Effect of Citalopram Pretreatment on Neuronal Responses to Neuropsychological Tasks in Normal Volunteers: An fMRI Study. *Neuropsychopharmacology*, *30*(9), 1724–1734. https://doi.org/10.1038/sj.npp.1300728

Dillo, W., Göke, A., Prox-Vagedes, V., Szycik, G. R., Roy, M., Donnerstag, F., … Ohlmeier, M. D. (2010). Neuronal correlates of ADHD in adults with evidence for compensation strategies – a functional MRI study with a Go/No-Go paradigm. *German Medical Science*, *8*, Doc09. https://doi.org/10.3205/000098

Horn, N. R., Dolan, M., Elliott, R., Deakin, J. F. W., & Woodruff, P. W. R. (2003). Response inhibition and impulsivity: an fMRI study. *Neuropsychologia*, *41*(14), 1959–1966. https://doi.org/10.1016/S0028-3932(03)00077-0

Liu, G.-C., Yen, J.-Y., Chen, C.-Y., Yen, C.-F., Chen, C.-S., Lin, W.-C., & Ko, C.-H. (2014). Brain activation for response inhibition under gaming cue distraction in internet gaming disorder. *The Kaohsiung Journal of Medical Sciences*, *30*(1), 43–51. https://doi.org/10.1016/j.kjms.2013.08.005

Maguire, R. P., Broerse, A., de Jong, B. M., Cornelissen, F. W., Meiners, L. C., Leenders, K. L., & den Boer, J. A. (2003). Evidence of enhancement of spatial attention during inhibition of a visuo-motor response. *NeuroImage*, *20*(2), 1339–1345. https://doi.org/10.1016/S1053-8119(03)00402-6

Menon, V., Adleman, N. E., White, C. D., Glover, G. H., & Reiss, A. L. (2001). Error-related brain activation during a Go/NoGo response inhibition task. *Human Brain Mapping*, *12*, 131–143. https://doi.org/10.1002/1097-0193(200103)12:33.0.CO;2-C

Mobbs, D., Eckert, M. A., Mills, D., Korenberg, J., Bellugi, U., Galaburda, A. M., & Reiss, A. L. (2007). Frontostriatal Dysfunction During Response Inhibition in Williams Syndrome. *Biological Psychiatry*, *62*(3), 256–261. https://doi.org/10.1016/j.biopsych.2006.05.041

Passamonti, L., Fera, F., Magariello, A., Cerasa, A., Gioia, M. C., Muglia, M., … Quattrone, A. (2006). Monoamine Oxidase-A Genetic Variations Influence Brain Activity Associated with Inhibitory Control: New Insight into the Neural Correlates of Impulsivity. *Biological Psychiatry*, *59*(4), 334–340. https://doi.org/10.1016/j.biopsych.2005.07.027

Penfold, C., Vizueta, N., Townsend, J. D., Bookheimer, S. Y., & Altshuler, L. L. (2015). Frontal lobe hypoactivation in medication-free adults with bipolar II depression during response inhibition. *Psychiatry Research: Neuroimaging*, *231*(3), 202–209. https://doi.org/10.1016/j.pscychresns.2014.11.005

Pornpattananangkul, N., Hariri, A. R., Harada, T., Mano, Y., Komeda, H., Parrish, T. B., … Chiao, J. Y. (2016). Cultural influences on neural basis of inhibitory control. *NeuroImage*, *139*, 114–126. https://doi.org/10.1016/j.neuroimage.2016.05.061

Shafritz, K. M., Bregman, J. D., Ikuta, T., & Szeszko, P. R. (2015). Neural systems mediating decision-making and response inhibition for social and nonsocial stimuli in autism. *Progress in Neuro-Psychopharmacology and Biological Psychiatry*, *60*, 112–120. https://doi.org/10.1016/j.pnpbp.2015.03.001

Stokes, P. R. A., Rhodes, R. A., Grasby, P. M., & Mehta, M. A. (2010). The Effects of The COMT val108/158met Polymorphism on BOLD Activation During Working Memory, Planning, and Response Inhibition: A Role for The Posterior Cingulate Cortex? *Neuropsychopharmacology*, *36*(4), 763–771. https://doi.org/10.1038/npp.2010.210

Townsend, J. D., Bookheimer, S. Y., Foland-Ross, L. C., Moody, T. D., Eisenberger, N. I., Fischer, J. S., … Altshuler, L. L. (2012). Deficits in inferior frontal cortex activation in euthymic bipolar disorder patients during a response inhibition task. *Bipolar Disorders*, *14*(4), 442–450. https://doi.org/10.1111/j.1399-5618.2012.01020.x

Völlm, B., Richardson, P., McKie, S., Elliott, R., Deakin, J. F. W., & Anderson, I. M. (2006). Serotonergic modulation of neuronal responses to behavioural inhibition and reinforcing stimuli: an fMRI study in healthy volunteers. *European Journal of Neuroscience*, *23*(2), 552–560. https://doi.org/10.1111/j.1460-9568.2005.04571.x

Völlm, B., Richardson, P., Stirling, J., Elliott, R., Dolan, M., Chaudhry, I., … Deakin, B. (2004). Neurobiological substrates of antisocial and borderline personality disorder: preliminary results of a functional fMRI study. *Criminal Behaviour and Mental Health*, *14*(1), 39–54. https://doi.org/10.1002/cbm.559
